# Identification of novel cellular intermediates unveils unique enzymes for flagellar glycan biosynthesis in *Clostridioides difficile*

**DOI:** 10.64898/2026.02.12.704994

**Authors:** Paul J. Hensbergen, Bob van Puffelen, August N.A. Ammerlaan, Paul L.C. Zuidgeest, Loes van Huijkelom, Rembrandt J.V. Kanbier, Maaike M. Vet, Xuan S. Zheng, Pranav V.N. Bhamidipati, Martin F. Larralde, Niek Blomberg, Sarantos Kostidis, Peter A. van Veelen, Robert A. Cordfunke, Jordy van Angeren, Arnoud H. de Ru, Zachary W.B. Armstrong, Wiep Klaas Smits, Dimitri V. Filippov, Martin Giera, Jeroen D.C. Codée, Jeroen Corver

**Affiliations:** Center for Proteomics and Metabolomics, Leiden University Medical Center, Leiden, 2333 ZA, The Netherlands; Leiden Institute of Chemistry, Leiden University, Leiden, 2333 CC, The Netherlands; Leiden University Center for Infectious Diseases, Leiden University Medical Center, Leiden, 2333 ZA, The Netherlands; Department of Immunology, Leiden University Medical Center, Leiden, 2333 ZA, The Netherlands

**Keywords:** flagella, glycosylation, bacteria

## Abstract

Glycosylation of bacterial surface proteins, such as flagellin (FliC), is important for their function and is often involved in virulence of pathogens. Glycans can be further modified by so-called post-glycosylation modifications (PGMs) often resulting in exclusive molecular structures. In *Clostridioides difficile* a unique glycan structure (Type A) decorates FliC (which forms the flagellar filament) that consists of an *O*-linked *N*-acetyl-β-D-glucosamine (GlcNAc) modified with an *N*-methyl-L-threonine via a phosphodiester linkage. This PGM is synthesized by a set of four enzymes encoded in one operon (*ftaABCD*), but the exact biosynthesis pathway and biosynthetic intermediates remain unknown.

In this study, we chemically synthesized two hitherto undescribed biosynthetic intermediates that we predicted based on bioinformatic analyses, CDP-threonine and CDP-*N*-methylthreonine. We showed that they are involved in the Type A PGM biosynthesis, as evidenced by mass spectrometric analyses of extracts of a set of *C. difficile* mutant strains. Furthermore, we characterized FtaC to be a SAM-dependent CDP-threonine *N*-methyltransferase, that installs the methyl group on CDP-threonine prior to transfer of the PGM to GlcNAc-FliC, and we revealed FtaD as the CDP-*N*-methylthreonine:GlcNAc *N*-methylthreoninephosphotransferase. Finally, using recombinantly expressed FtaC and FtaD in combination with synthetic CDP-threonine, we reconstituted the biosynthesis pathway of the Type A PGM *in vitro*.

Overall, our results open avenues to explore these unique biosynthesis enzymes in molecular detail to provide new points of entry for the development of biosynthesis inhibitors and tools to study the role of this PGM in virulence and flagellar assembly.

## Introduction

Glycosylation is a pivotal process in all domains of life. In bacteria, structures containing glycans are important virulence mediators, not only in the form of glycoconjugates such as lipopolysaccharides and capsular polysaccharides, but also as glycoproteins (1, 2). Glycosyltransferases are the main enzymes involved in protein glycosylation which use nucleotide-sugars, e.g. UDP-GlcNAc/GalNAc, GDP-mannose/fucose and CMP-sialic acid, as donor substrates (3, 4). The diversity of glycan structures can be further expanded by additional modifications such as methylation, acetylation, phosphorylation and sulfation, collectively known as post-glycosylation modifications (PGMs) (5, 6). Due to rare structural elements, including unique PGMs (7), and unusual glycosylation patterns, bacterial glycosylation provides promising targets for new, highly-specific, treatment strategies (8).

Most bacterial glycoproteins are found on cell-envelope structures (9-12). One such glycoprotein is flagellin (FliC), that polymerizes to form a major part of the flagellar filament (13, 14). A multitude of flagellar glycan structures have been described, which differ between and within species (15-21). Beyond their role in motility (22, 23), flagella are critical factors in bacterial pathogenesis (24, 25). This is the case in *Clostridioides difficile* (26), a notorious gut pathogen (27) that can cause symptoms that vary from mild diarrhea to life-threatening pseudomembranous colitis and toxic megacolon (28). Notably, in *C. difficile* 630Δ*erm* FliC is decorated with a unique flagellar glycan structure (Type A) consisting of an *O*-linked β-*N*-acetylglucosamine (GlcNAc) that is modified with an *N*-methyl-L-threonine via a phosphodiester linkage at the C-3 position ((29), **Figure 1A**). Of note, this flagellar glycan impacts motility, cell surface properties and virulence (29). A set of five genes is linked to the biosynthesis of the flagellar Type A glycan (**Figure 1B**). The glycosyltransferase FgtA (**F**lagellin **G**lycosyl**T**ransferase **A**), encoded by *cd0240* in *C. difficile* 630Δ*erm* is pivotal for FliC glycosylation and disruption of this gene leads to non-glycosylated FliC (30). The modification of the GlcNAc with the *N*-methyl-phosphothreonine in *C. difficile* 630Δ*erm* is dependent on the *cd0241*-*cd0244* operon (31), but the nature of the enzymatic steps, and the role and structure of potential biosynthetic intermediates, remains largely unknown. We will refer to products of these genes as Fta (<u>F</u>lagellin <u>T</u>ype <u>A</u>) ABCD (**Figure 1B**). Building upon bioinformatics analyses and mass spectrometry-based proteomics data of all four individual *C. difficile* 630*Δerm fta* mutants, we previously proposed a biosynthetic pathway for the Type A glycan (29, 31, 32). The crucial component in the model (**Figure 1C**) is the so far undescribed high-energy intermediate cytidine-5’-diphosphate-activated threonine (CDP-threonine, CDP-Thr) and its *N*-methylated counterpart (CDP-*N*-methylthreonine, CDP-Me-Thr). Although CDP-activated conjugates are well known intermediates for the synthesis of phospholipids (33) (e.g. CDP-choline and CDP-ethanolamine), bacterial teichoic acids (CDP-ribitol and CDP-glycerol (34)), and α-dystroglycan glycans (CDP-ribitol (35, 36)), only a single example of a CDP-amino acid intermediate has hitherto been described (37). In our model, we have proposed that CDP-Thr is generated from cytidine triphosphate (CTP) and phosphothreonine, itself generated form phosphoserine. CDP-threonine is methylated, and subsequently the *N*-methylated phosphothreonine moiety is mounted on GlcNAc-FliC to complete the biosynthesis of the Type A glycan (**Figure 1C**). Although methylation is predicted to occur on CDP-threonine, the methyl group could also be transferred earlier or later in the pathway.

**Figure 1:**
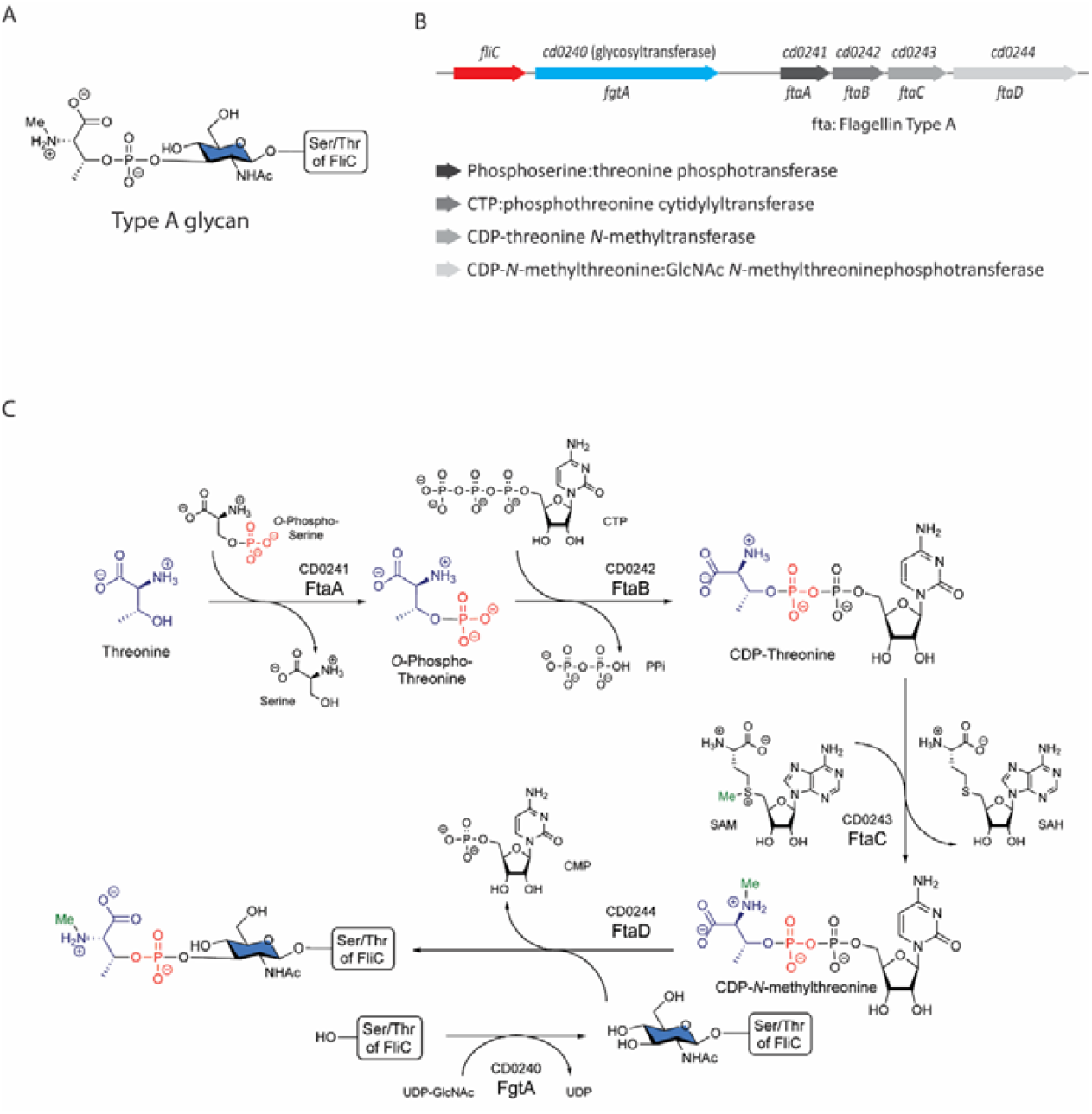
*C. difficile* flagellar Type A glycan: structure, gene cluster and biosynthesis model. **A**: Structure of the flagellar Type A glycan in C. *difficile* (29) **B**: Gene cluster involved in de biosynthesis of the Type A glycan on flagellin C (FliC), and (predicted) functions of the proteins encoded by fgtA/cd0240 (flagellin glycosyltransferase) and *ftaABCD* (*cd0241*-*cd0244*). **C**: Model showing the predicted individual steps of the Type A glycan biosynthesis. CTP/CDP/CMP: cytidine tri/di/monophosphate. PPi: pyrophosphate. UDP: uridine diphosphate. SAM: S-adenosylmethionine. SAH: *S*-adenosylhomocysteine.

In this study, we report on the discovery of CDP-Thr and CDP-Me-Thr as novel cellular metabolites and as substrates for two unique enzymes in the Type A glycan biosynthesis pathway in *C. difficile*. We have generated synthetic CDP-Thr and CDP-Me-Thr, which we used as standards for metabolomics analyses of *C. difficile* extracts and as substrates for recombinantly expressed and purified FtaC and FtaD. Using a series of *C. difficile* mutant strains we show how the presence of CDP-Thr and CDP-Me-Thr is in line with their role as intermediates of the Type A glycan biosynthesis. We further show that FtaC and FtaD have CDP-threonine *N*-methyltransferase and CDP-*N*-methylthreonine:GlcNAc N-methylthreoninephosphotransferase activity, respectively, and show that they act in concert, recapitulating the last two steps of the Type A PGM in vitro. Our results offer insight into the role of novel CDP-conjugated cellular metabolites in the biosynthesis of a glycan structure on flagellin and provide the foundation for studying similar pathways in other pathogens, such as *Pseudomonas aeruginosa*.

## Results

### Synthesis, characterization and HILIC-MS detection of CDP-threonine and CDP-N-methylthreonine

In our model for the biosynthesis of the Type A glycan in *C. difficile* (**Figure 1**), we predict two critical molecular intermediates that have so far remained elusive: CDP-threonine (CDP-Thr) and CDP-*N-* methylthreonine (CDP-Me-Thr). To accomplish their unambiguous identification, we undertook the *de novo* organic synthesis of both molecules (**Figure 2A, Supplemental Figure S1**). Pyrophosphates are energy-rich metabolites and intrinsically labile, owing to the high reactivity of the pyrophosphate bond. In addition, the presence of the pyrophosphate functionality on the β-carbon of threonine renders the CDP-(*N-*methyl)threonine molecules susceptible to β-elimination. To generate CDP-Thr and CDP-Me-Thr we turned to chemistry we previously developed for the synthesis of nucleotide diphosphate sugar donors, which hinges on the coupling of a phosphomonoester and a phosphoramidite (38-40). Phosphoramidites are amongst the most effective phosphorylating agents and they can be activated under mild conditions to generate a strong electrophile that can react with the phosphomonoester to provide a P(III)-P(V) intermediate. Subsequent *in situ* oxidation then provides the (partially protected) pyrophosphate. Here we chose to equip the threonine building block with the phosphomonoester functionality and couple it with a suitably protected cytidine phosphoramidite. Because of the lability of the target compounds towards nucleophilic or basic conditions, we implemented a protecting group strategy based on reductively cleavable and acid-labile masking groups. Thus, the carboxylic acid function in *N*-Boc threonine **1** was masked as a *tert*-butyl ester, after which the phosphate was installed. To this end, di(9-fluorenylmethyl)-*N,N*-diisopropylphosphoramidite was activated with dicyanoimidazole (DCI) as activator and coupled to the threonine hydroxyl. Subsequent oxidation of the so-formed phosphite intermediate and mild basic deprotection of the 9-fluorenylmethyl groups delivered the suitably protected threoninephosphate **3**. In parallel, the required cytidine phosphoramidite **5** was assembled. We equipped this building block with benzyloxycarbonyl (Cbz) protecting groups, to endow the upcoming synthetic intermediates with lipophilic character to enable HPLC purification. The building block was generated by silylation of the primary alcohol of cytidine **4**, installation of the Cbz-groups on the remaining alcohols and exocyclic cytosine nitrogen, removal of the silyl ether and installation of the phosphoramidite. The union of threonine phosphate **3** and cytidine phosphoramidite **5** was effected under the agency of DCI, after which the P(V)-P(III) intermediate was oxidized and the cyanoethyl group was removed. For purification purposes the *tert*-butyl ester and carbamate were unmasked to provide the semi-protected pyrophosphate **6**, that was purified by HPLC. Final reductive removal delivered the target CDP-threonine, that was purified using size-exclusion chromatography. Here it was noted that a mildly acidic eluent (0.15 M NH_4_OAc in water) was required, as the use of dilute NH_4_OAc (pH ∼ 7) led to hydrolysis of the pyrophosphate during concentration, forming threonine phosphate and cytidine phosphate, indicating the lability of the metabolite. Using a similar approach we generated CDP-Me-Thr (See **Supplemental data chemistry** for details). Both building blocks were fully characterized using ^1^H-^13^C, ^31^P -NMR and mass spectrometry and obtained in multi-mg amounts to allow for their use as standards in our metabolomics assays, but also to reconstitute the biosynthesis pathway on scale. For the projected metabolomics studies, we set-up a negative ion mode HILIC-MS workflow for the analysis of both molecules. Using this platform, CDP-Thr and CDP-Me-Thr could be resolved with baseline separation (**Figure 2B**).

**Figure 2:**
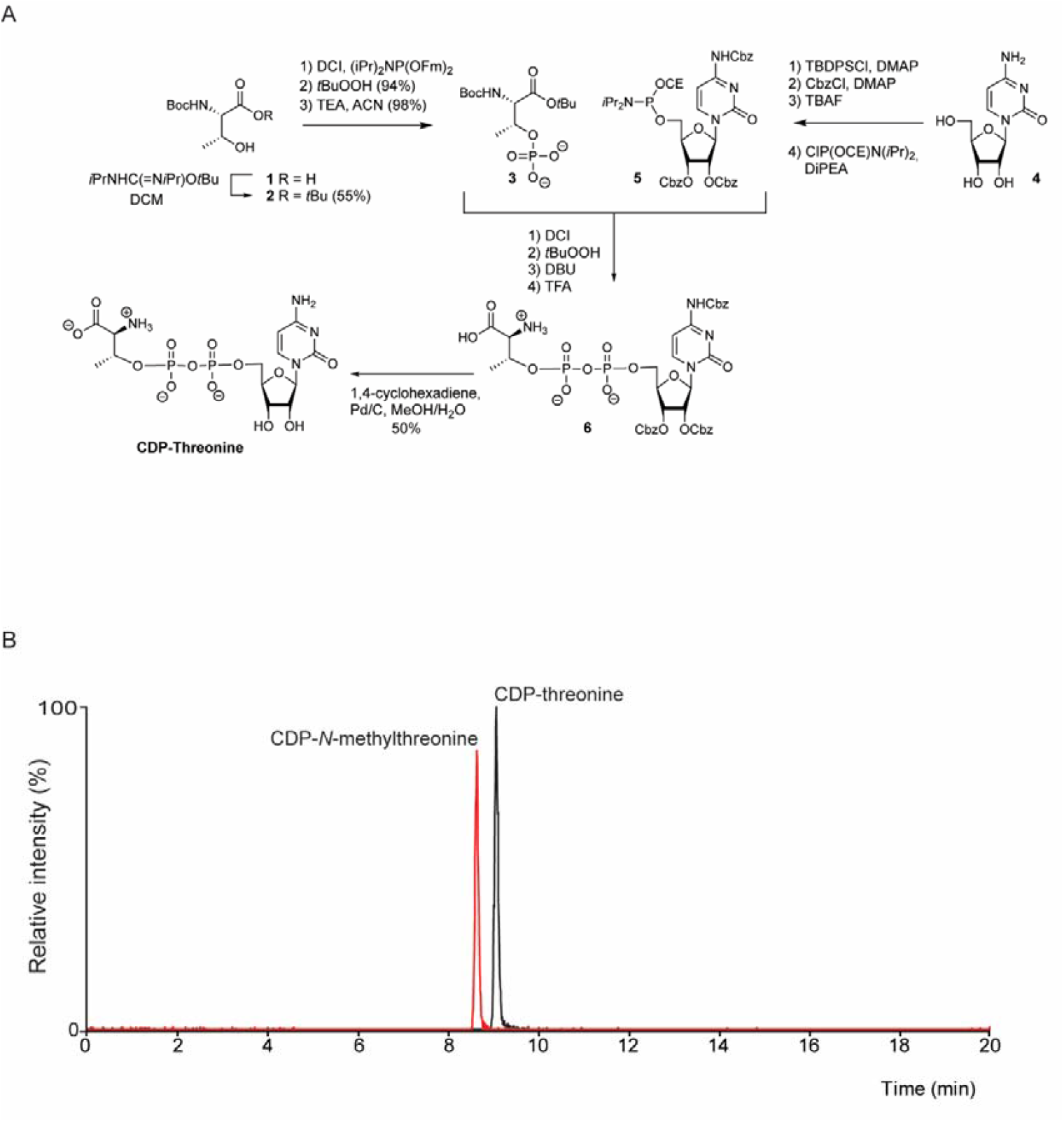
Synthesis, characterization and HILIC-MS detection of CDP-threonine (CDP-Thr) and CDP-*N*-methylthreonine (CDP-Me-Thr) **A:** Synthesis of CDP-Thr. For synthesis of CDP-*N*-methylthreonine, see Supplemental Figure S1. For full details see Supplemental data chemistry. **B:** An equimolar amount of CDP-Thr and CDP-Me-Thr were analysed by HILIC-MS. Shown are the extracted ion chromatograms of CDP-Thr (black line, [M-H]^-^, theoretical *m/z* 503.059) and CDP-Me-Thr (red line, [M-H]^-^, theoretical *m/z* 517.074).

### CDP-*N*-methylthreonine is a cellular metabolite in C. *difficile* strains with the FliC Type A glycan

To determine whether CDP-Me-Thr is present in *C. difficile* strain 630Δ*erm*, which is known to display the Type A glycan on its flagella (29-31), we analyzed an extract of these cells on our HILIC-MS platform. Notably, we observed a clear signal corresponding to CDP-Me-Thr (**Figure 3A**) at the same retention time as our analytical standard (**Figure 3B**). Furthermore, MS/MS fragmentation patterns of the synthetic and the endogenous CDP-Me-Thr were in excellent agreement (Supplemental Figure S2A). As a control, we also performed these experiments with the *C. difficile* strain R20291 (RT027).

**Figure 3:**
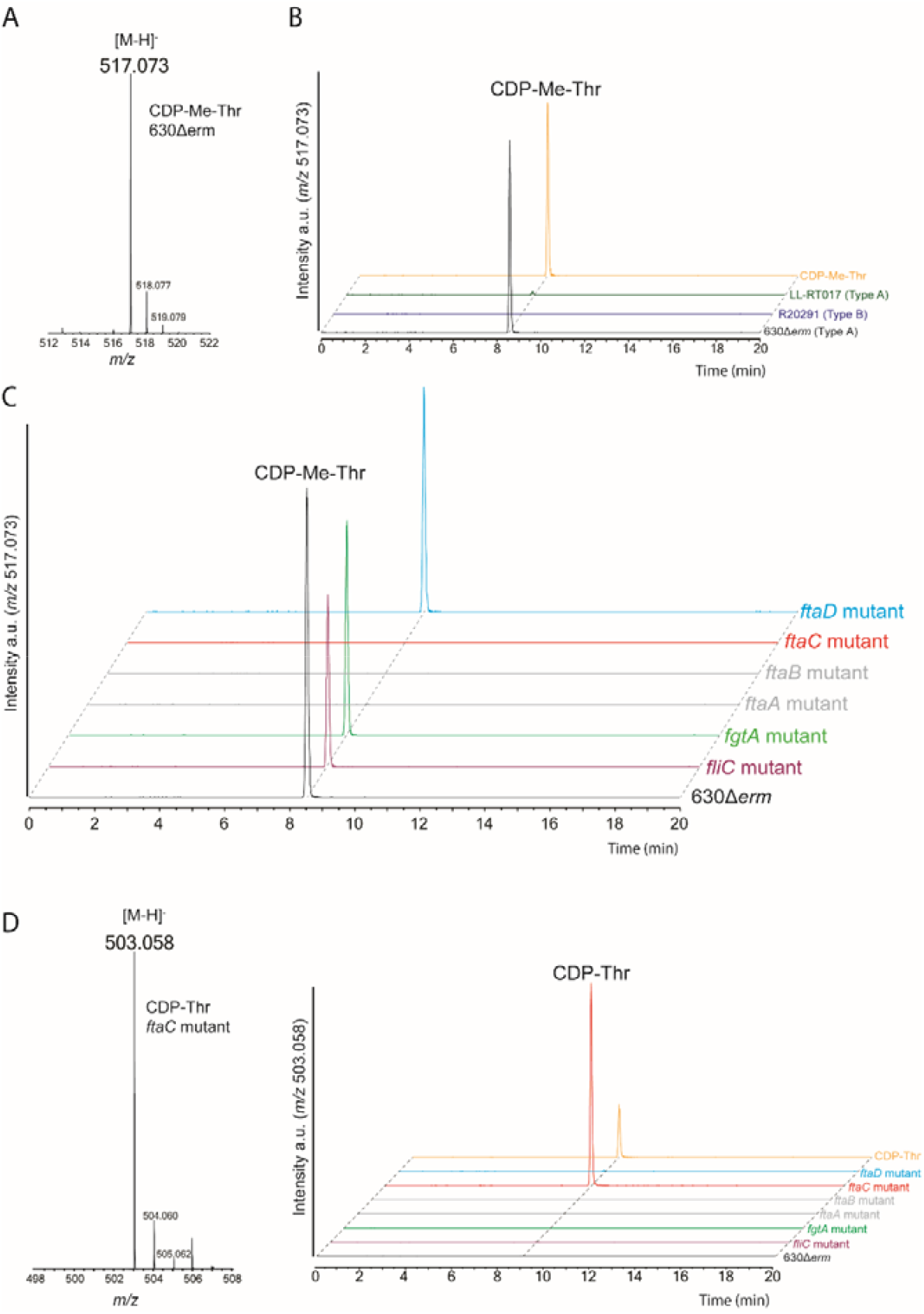
CDP-*N*-methylthreonine (CDP-Me-Thr) and CDP-threonine (CDP-Thr) are novel cellular intermediates which are altered in individual Type A biosynthetic *C. difficile* gene mutants. **A**: HILIC-MS identification of CDP-Me-Thr ([M-H]^-^, theoretical *m/z* 517.074) in an extract of *C. difficile* strain 630Δ*erm*. See Supplemental Figure S2A for the corresponding MS/MS spectrum. **B:** HILIC-MS analysis of synthetic CDP-Me-Thr (standard) and extracts of *C. difficile* strains 630Δ*erm* (Type A), R20291 (Type B) and LL-RT017 (Type A). Shown are the extracted ion chromatograms of *m/z* 517.073, corresponding to CDP-Me-Thr. **C:** HILIC-MS analysis of extracts of WT *C. difficile* strain 630Δ*erm* and fgtA, *ftaA*-*ftaD* and *fliC* mutant strains. Shown are the extracted ion chromatograms of *m/z* 517.073, corresponding to CDP-Me-Thr. **D:** HILIC-MS analysis of synthetic CDP-Thr (standard) and extracts of WT *C. difficile* strain 630Δ*erm* and fgtA, *ftaA*-*ftaD* and *fliC* mutant strains. Shown are the extracted ion chromatograms of *m/z* 503.058, corresponding to CDP-Thr (MS signal in *ftaC* mutant shown in the left panel, see Supplemental Figure S2B for the corresponding MS/MS spectrum).

This strain contains a different glycan modification, Type B, whose synthesis does not involve CDP-Me-Thr. Indeed, CDP-Me-Thr was not observed in strain R20291 (RT027) (**Figure 3B**). The Type A biosynthesis gene cluster is also present in PCR ribotype 017 strains (29). Using proteomics, we confirmed that FliC is modified with Type A glycans in a reference strain (LL-RT017, (41)) of this PCR ribotype (**Supplemental Figure S3A**). However, the intensity of FliC peptides was much lower as compared to the 630Δ*erm* strain (**Supplemental Figure S3B**). Consistent with this, the level of CDP-Me-Thr was much lower in the LL-RT017 strain (**Figure 3B**).

Collectively, these results showed the presence of the novel cellular metabolite CDP-Me-Thr in *C. difficile* strains with the FliC Type A glycan, but not in a strain with Type B glycans.

### CDP-N-methylthreonine is absent in C. *difficile* 630Δerm ftaA, ftaB and ftaC mutant strains

We next studied the presence of CDP-Me-Thr in individual *ftaA-ftaD* mutant strains of *C. difficile* 630*Δerm* (31). If the presence of CDP-Me-Thr were the result of the sequential activity of FtaA, FtaB and FtaC, we would expect it to be absent in the corresponding individual mutant strains. On the other hand, we expected the *ftaD* mutant strain to be competent in synthesizing CDP-Me-Thr as its predicted activity is to transfer the *N*-methyl-phosphothreonine to the GlcNAc (**Figure 1C**). We analyzed extracts of *ftaA-ftaD* mutant strains by HILIC-MS. In accordance with our model, our results showed that, in addition to WT, CDP-Me-Thr was observed in *ftaD* mutant cells, but not in *ftaA, ftaB* and *ftaC* mutant cells (**Figure 3C**).

Given that the *O*-GlcNAcylation of FliC occurs in parallel to the synthesis of CDP-Me-Thr (**Figure 1C**), we also expected to observe CDP-Me-Thr in a *fgtA* and *fliC* mutant strain. Indeed, HILIC-MS analysis of lysates of these mutants clearly demonstrated the presence of CDP-Me-Thr in these cells (**Figure 3C**).

### CDP-threonine accumulates in C. *difficile* 630Δerm ftaC mutant cells

To validate our prediction that the methylation of CDP-Thr is a discrete step in the Type A biosynthetic pathway (**Figure 1C**), we assessed the presence or absence of CDP-Thr in a wild-type strain and in the panel of mutant strains described above. We could clearly demonstrate the presence of CDP-Thr in *ftaC* mutant cells (**Figure 3D, Supplemental Figure S2B**), but not in *ftaA* and *ftaB* mutant cells (**Figure 3D**). Only a trace of CDP-Thr was observed in one of the other extracts (*ftaD* mutant cells). These results demonstrate that CDP-Thr is an intermediate of the Type A biosynthesis and the absence of FtaC results in the accumulation of CDP-Thr.

Overall, our metabolite analyses demonstrate for the first time that both CDP-Thr and CDP-Me-Thr are present in vivo and provide evidence for a role of these molecules as intermediates of the flagellar Type A glycan biosynthesis in *C. difficile*. Moreover, in line with our model (**Figure 1C**), these data support a role for FtaA and FtaB in the synthesis of CDP-Thr, for FtaC in the methylation of CDP-Thr, and for FtaD to complete the Type A biosynthesis.

### FtaC is a CDP-threonine N-methyltransferase

In an *ftaC* mutant strain, FliC is modified with a Type A structure lacking the methyl group (29, 31) in-line with the predicted methyltransferase activity of FtaC (**Figure 1B**). Our metabolomic analyses showed a high level of CDP-Thr only in *ftaC* mutant cells (**Figure 3D**). This indicates that CDP-Thr is a direct substrate of FtaC (**Figure 1C**). To test this, we performed *in vitro* experiments with recombinant His6-FtaC (**Supplemental Figure S4**) and the universal methyl donor *S*-adenosylmethionine (SAM). Upon methyl transfer, SAM is converted to S-adenosylhomocysteine (SAH), which can be monitored in a bioluminescent-based assay (**Figure 4A**). It was found that incubation of FtaC with CDP-Thr and SAM, but not CDP-Thr or SAM alone, resulted in a SAH formation (**Figure 4B**). No activity of FtaC was observed when threonine or phosphothreonine were used as potential methyl acceptors (**Figure 4B**).

**Figure 4:**
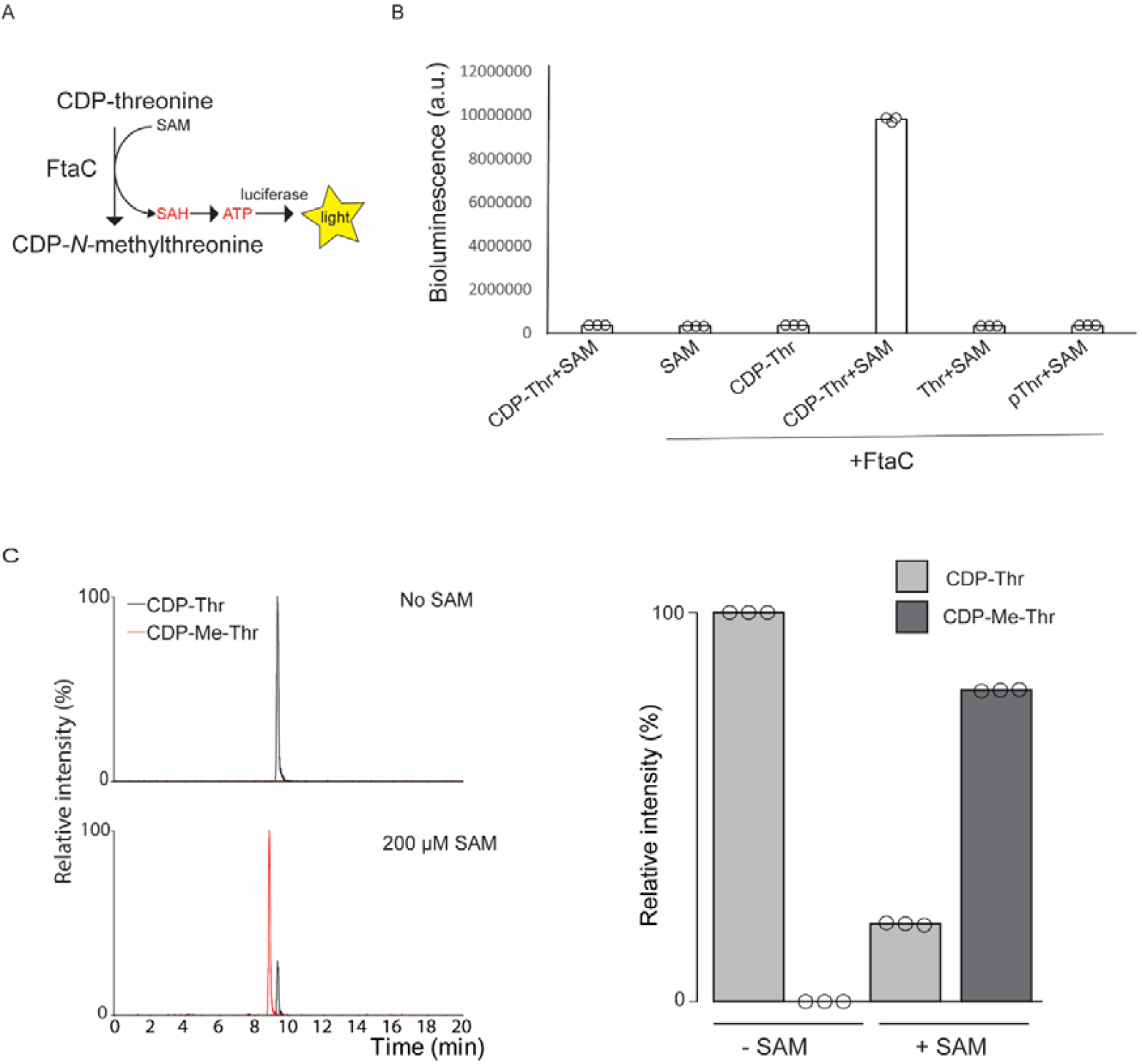
FtaC is a (SAM-dependent) CDP-threonine *N*-methyltransferase. **A:** Schematic representation of the bioluminescent-based methyltransferase assay (MTase-Glo™ assay) for the monitoring of FtaC activity. SAM: *S*-adenosylmethionine. SAH: *S*-adenosylhomocysteine. ATP: adenosine triphosphate. **B:** FtaC was incubated with different (combinations of) donor and acceptor substrates (CDP-Thr, pThr and Thr at 100 µM, SAM at 200 µM) and activity was subsequently detected using the bioluminescent-based methyltransferase assay (**A**). An assay with CDP-Thr+SAM without FtaC was used as a negative control. **C**: FtaC was incubated with CDP-Thr (100 µM) with and without SAM (200 µM) and subsequently analyzed by HILIC-MS. Left panel: Extracted ion chromatograms of *m/z* 503.059 and 517.074, corresponding to CDP-Thr and CDP-Me-Thr respectively. Right panel: relative abundance of CDP-Thr (substrate) and CDP-Me-Thr (product) in triplicate FtaC experiments using 100 µM CDP-Thr with and without SAM, respectively.

To verify the product of FtaC activity was indeed CDP-Me-Thr, we analyzed the products of the *in vitro* reactions using HILIC-MS (**Figure 4C**, left panel). The results showed reproducible conversion of CDP-Thr to CDP-Me-Thr (**Figure 4C**, right panel) in the presence of FtaC and SAM. Although di- and trimethylation have been observed with other methyltransferases (42, 43), we found no evidence for these products in our data, indicating that CDP-Me-Thr is most likely the only product of the FtaC-mediated reaction.

Overall, these data firmly establish FtaC as an SAM-dependent CDP-threonine *N*-methyltransferase.

### FtaD is a phosphotransferase that installs the Type A PGM on a O-GlcNAc peptide, using CDP-Me-Thr as the donor substrate

We have previously shown that in *ftaD* mutant cells FliC is only modified with a *N*-acetylglucosamine residue (31). Given the putative phosphotransferase activity of FtaD, we predict that FtaD catalyzes the transfer of the *N-*methyl-phosphothreonine onto the GlcNAc, using CDP-Me-Thr as the donor substrate (**Figure 1C**). To test this, we performed *in vitro* experiments with recombinant His6-FtaD (**Supplemental Figure S4**), CDP-Me-Thr and a FliC-derived O-GlcNAc peptide (TMVsSLDAALK, s=Ser-β-D-GlcNAc) (31) as the acceptor substrate. Notably, LC-MS/MS analysis showed FtaD-dependent transfer of the *N*-methyl-phosphothreonine to the GlcNAc, resulting in a full Type A structure on the peptide (**Figure 5A**). The fragmentation spectrum of the product peptide (**Supplemental Figure S5**) showed characteristic fragment ions at *m/z* 214.048 and 284.053 (**Supplemental Figure 3A**), that are well-known from endogenous Type A-modified peptides (44), confirming the phosphodiester linkage between the GlcNAc and the *N*-methylthreonine.

**Figure 5:**
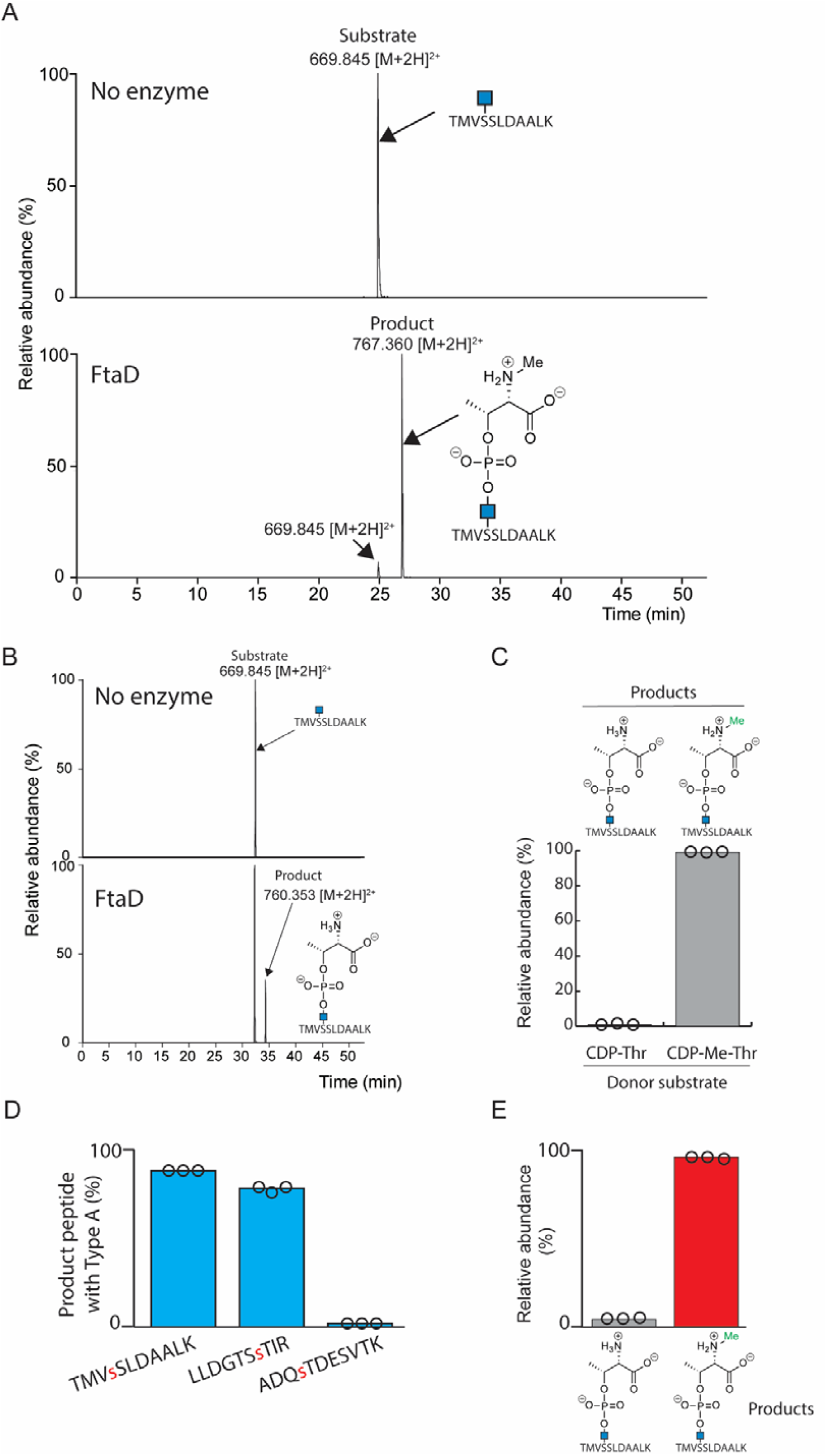
FtaD is a CDP-*N*-methylthreonine:GlcNAc *N*-methylthreoninephosphotransferase. **A**: FtaD was incubated with CDP-Me-Thr and an acceptor substrate peptide containing an *O*-GlcNAc (TMVsSLDAALK, s=Ser-β-D-GlcNAc). Product formation was detected using LC-MS/MS. Shown are the extracted ion chromatograms for the acceptor substrate peptide (*m/z* 669.845, [M+2H]2^+^) and the product peptide with the full Type A structure (*m/z* 767.360, [M+2H]^2+^). The MS/MS spectra of both peptides can be found in Supplemental Figure S5. An incubation without enzyme was used as a control. **B**: FtaD was incubated with CDP-Thr and an acceptor substrate peptide containing an *O*-GlcNAc (TMVsSLDAALK, s=Ser-β-D-GlcNAc). Product formation was detected using LC-MS/MS. Shown are the extracted ion chromatograms for the acceptor substrate peptide (*m/z* 669.845, [M+2H]2^+^) and the product peptide with the Type A structure lacking the N-methyl moiety (*m/z* 760.353, [M+2H]^2+^). An incubation without enzyme was used as a control. **C**: FtaD was incubated with an equimolar mixture of CDP-Thr CDP-Me-Thr, and an acceptor substrate peptide containing an *O*-GlcNAc (TMVsSLDAALK, s=Ser-β-D-GlcNAc). Product formation was detected using LC-MS/MS. Shown are the relative abundances of the product peptide with the Type A structure lacking the N-methyl moiety (*m/z* 760.353, [M+2H]2^+^) and the full Type A structure (*m/z* 767.360, [M+2H]2^+^), respectively. **D**: FtaD was incubated with CDP-Me-Thr and three different acceptor peptides containing an *O*-GlcNAc (s=Ser-β-D-GlcNAc). LC-MS/MS was then performed to determine the relative abundances of the product peptides (as percentage of the sum of the intensities of the product and substrate peptides) **E**: FtaC and FtaD were incubated with CDP-Thr, SAM and an acceptor substrate peptide containing an *O*-GlcNAc (TMVsSLDAALK, s=Ser-β-D-GlcNAc). Product formation was detected using LC-MS/MS. Shown are the relative abundances of the product peptide with the Type A structure lacking the N-methyl moiety (*m/z* 760.353, [M+2H]^2+^) and the full Type A structure (*m/z* 767.360, [M+2H]^2+^), respectively.

We previously observed that FliC was modified with a Type A structure lacking the methyl group in an *ftaC* mutant (31). In these cells, we also observed the accumulation of CDP-Thr (**Figure 3D**), suggesting that FtaD can (also) use CDP-Thr as the donor substrate. To investigate this *in vitro*, we performed the transfer assay with FtaD and CDP-Thr as the donor substrate. The results from this experiment demonstrated the formation of a peptide with the Type A glycan lacking the methyl group (**Figure 5B**). However, based on the relative intensity of the product peptide, the efficiency of the reaction appeared to be significantly less than the reaction using CDP-Me-Thr as the donor substrate (**Figure 5A**). To assess a difference in efficiency of the transfer reaction with both donor substrates, we performed experiments with an equimolar amount of CDP-Thr and CDP-Me-Thr as substrates in a single reaction. We found ∼100-fold more product peptide with the full Type A compared to the structure lacking the methyl group, demonstrating that FtaD preferentially uses CDP-Me-Thr over CDP-Thr as a donor substrate (**Figure 5C**).

In *C. difficile* 630*Δerm*, FliC is modified with the Type A glycan at many different sites (30). To demonstrate the specificity of the FtaD-mediated transfer reaction, we also performed our assay using another FliC-derived peptide (LLDGTSsTIR, s=Ser-β-D-GlcNAc) and a synthetic peptide unrelated to FliC, but also carrying a *O*-GlcNAc (ADQsTDESVTK (s=Ser-β-D-GlcNAc). The results also showed transfer of the *N*-methyl-phosphothreonine onto the FliC-derived peptide, with an efficiency comparable to that of the TMVsSLDAALK acceptor peptide, but not to the synthetic peptide unrelated to FliC (**Figure 5D**). Because in FliC some Type A-modified residues are very close together, we also tested an acceptor peptide bearing two *O-*GlcNAcs, mimicking two sites in FliC. With this peptide a mixture of products with either one or two Type A structures was observed (**Supplemental Figure S6**), indicating that FtaD can transfer the *N*-methyl-phosphothreonine to multiple sites on the same peptide.

Overall, the above data establish FtaD as a genuine CDP-*N*-methylthreonine:GlcNAc *N*-methylthreoninephosphotransferase.

### FtaC and FtaD act in concert to modify an *O*-GlcNAc-peptide *in vitro*

Having characterized the enzymatic activity of the individual proteins FtaC (**Figure 4**) and FtaD (**Figure 5**), we sought to recapitulate the Type A biosynthetic pathway starting from CDP-threonine *in vitro* as the CDP-Me-Thr generated by FtaC is believed to be used by FtaD to modify FliC in vivo (**Figure 1C and 4**). Thus, we incubated CDP-Thr, SAM and a FliC model *O*-GlcNAc peptide (TMVsSLDAALK, s=Ser-β-D-GlcNAc) with FtaC and FtaD. Subsequent LC-MS/MS analyses of this coupled assay unambiguously showed the formation of the FliC-derived peptide with a full Type A structure (**Figure 5E**). These results directly demonstrate that the CDP-Me-Thr produced by FtaC can be used by FtaD.

Collectively, the above data firmly establish the role of FtaC and FtaD in the biosynthesis of the unique Type A PGM in *C. difficile*.

### *ftaABCD* homologs are found in other bacterial species

The enzymatic activities we characterized are likely present in different bacterial species. As a notable example, a similar FliC glycan structure is observed in *Pseudomonas aeruginosa*, where it is called b-type (45). Interestingly, in contrast to *C. difficile*, the glycosyltransferase and phosphotransferase activities are fused into one protein (PA1091) in *P. aeruginosa* (31). Extensive analysis of other sequenced bacterial genomes revealed the presence of several species that contained either a *C. difficile* Type A-like gene cluster or a *P. aeruginosa* b-type-like gene cluster (**Figure 6**) and many species that appear to encode some, but not all, enzymes involved in type A biosynthesis (**Supplemental table S1**).

**Figure 6:**
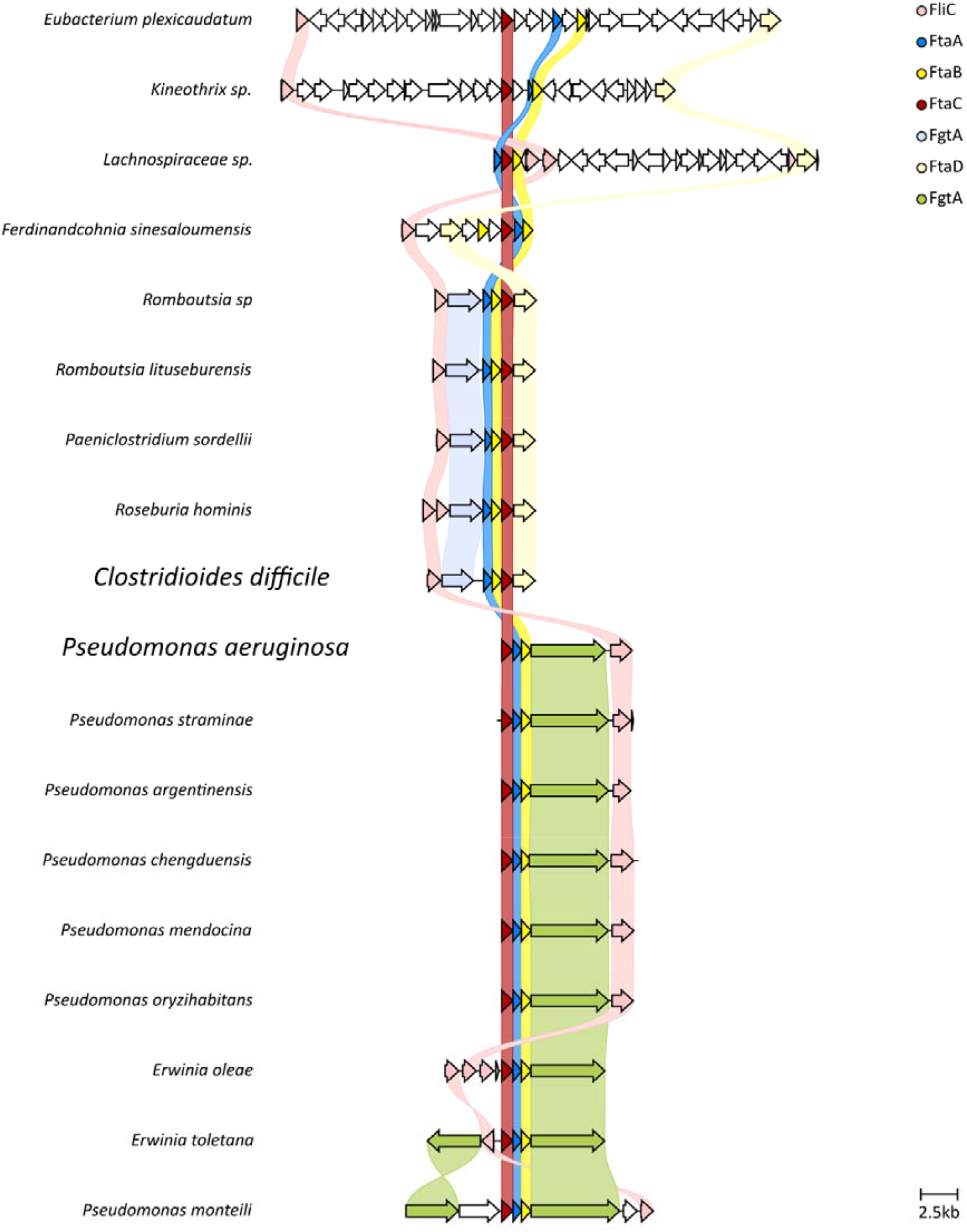
Overview of biosynthetic gene clusters encoding enzymes involved in type A (upper figure) and b-type (lower figure) flagellin glycan modifications. Type of enzymes are indicated by color and homologues are connected with lines of the same color. Colors are explained on the right. Grey color indicates genes that are not conserved between the different species.

*P. sordellii, R. hominis, Romboutsia sp*., and *R. lituseburensis* (all belonging to the phylum Bacillota) contain gene clusters highly similar to the *ftaABCD*-operon adjacent to the *fliC* gene, indicating that their flagella likely contain modifications very similar to Type A. In other species (*F. sinesaloumensis, E. caudatum, Kineothix sp and Lachnospiraceae sp*) genes with similarity to the *fta* operon appear to be part of a much larger cluster of genes. These clusters contain several additional glycosyltransferases and other genes, suggesting that the final glycan structure is more complex than Type A, but likely contains a *N*-methyl-phosphothreonine.

We found 5 species (*P. argentinensis, P. straminae, P. mendocina, P. chengduensis and P. oryzihabitans;* all belonging to the phylum Pseudomonadota) with gene clusters highly similar to the *P. aeruginosa pa1088-1091* operon (**Figure 6**). Three other species contained FliC-associated gene clusters that were more divergent, but nevertheless retained the organization of the enzymes putatively involved in the post-glycosylation modification. We expect that these species contain glycans that are modified with a PGM like that of the b-type glycan in *P. aeruginosa*.

## Discussion

Post-glycosylation modifications (PGMs) have a significant impact on the diversity of glycan structures and characterization of the underlying biosynthesis pathways often require unique metabolites and enzymatic activities. Specific PGMs involved in virulence of bacterial pathogens are particularly interesting because their biosynthesis pathways may provide novel strategies for therapeutic intervention (46). The Type A glycan structure in the human pathogen *C. difficile*, involved in virulence (29), is distinct because it contains a GlcNAc that is modified with an *N*-methylthreonine through a phosphodiester bond (29). Some other bacterial PGMs involving amino acids have been described (47-49) but these are linked via an amide bond and are probably independent of CDP-conjugated amino acids, although many of their biosynthesis pathways have not yet been fully elucidated.

We here, for the first time, identify CDP-activated threonine intermediates *in vivo* and demonstrate how the presence/absence in individual biosynthetic *C. difficile* gene mutant strains is consistent with their role in the Type A glycan biosynthesis pathway. Two enzymatic steps of this pathway were confirmed *in vitro*, thereby establishing novel SAM-dependent CDP-threonine *N*-methyltransferase (FtaC) and CDP-*N*-methylthreonine:GlcNAc *N*-methylthreoninephosphotransferase (FtaD) activities.

To our knowledge, CDP-glutamine is the only other CDP-conjugated amino acid that has hitherto been described. It acts as an intermediate in the biosynthesis of a *Campylobacter jejuni* capsular polysaccharide modification (37). However, in this pathway CDP-glutamine is hydrolyzed to cytidine diphosphoramidate and not used as a donor substrate for a phosphotransferase. Moreover, CDP-glutamine has only been studied *in vitro* and has not been described in *C. jejuni* cells to date.

In the present work, we showed that FtaC-generated CDP-Me-Thr acts as a donor for the phosphotransferase FtaD, which conjugates phospho-*N*-methylthreonine to a GlcNAc-functionalized FliC peptide, thereby expanding the catalytic repertoire of the EC 2.7.8 family of transferases (“Transferases for other substituted phosphate groups”, Nomenclature Committee of the International Union of Biochemistry and Molecular Biology (NC-IUBMB), https://iubmb.qmul.ac.uk/enzyme/). More specifically, FtaD is a new member of the CDP-alcohol phosphotransferase (CDP-AP) family which transfers substituted phosphate groups from a CDP-linked donor to an alcohol-acceptor. Notably, FtaD lacks the classical CDP-AP motif (50) which is known from enzymes in phospholipid biosynthesis (33). Other CDP-APs lacking this motif include TagF (51), TarL (52), Fukutin and Fukutin-related protein (53). Based on a BLASTP analysis, FtaD (UniprotID: Q18CY0) was previously found to have similarity to TagF (30), an enzyme that uses CDP-glycerol as a donor substrate for teichoic acid biosynthesis (51). A Foldseek analysis using the AlphaFold (v 2.0)-predicted structure (54) also revealed similarity of FtaD and the *Staphylococcus aureus* TagF-like protein TarF, which uses CDP-ribitol as the donor substrate. However, we here established FtaD as a unique phosphotransferase involved in the Type A glycan biosynthesis, arguing against a role for FtaD in cell surface polysaccharide biosynthesis. This is consistent with the fact that another phosphotransferase (CD2775) is involved in this process in *C. difficile* 630*Δerm* (55, 56).

The level of CDP-Me-Thr in the *ftaD* mutant strain was similar to that in WT cells, suggesting that it is regulated irrespective of whether CDP-Me-Thr is consumed as a donor substrate in Type A glycan biosynthesis. We additionally observed CDP-Me-Thr in strain LL-RT017, which also has the Type A glycan biosynthesis cluster, but the levels appeared much lower than in the 630Δ*erm* WT strain, which may relate to different expression levels of the flagellar operon (57, 58). As expected, we did not observe CDP-Me-Thr in *C. difficile* strain R20291, which harbors the Type B flagellar glycan. In contrast to Type A, the *C. difficile* Type B glycan contains multiple monosaccharides (59) and only the GlcNAc is common between both glycan structures in *C. difficile*.

We demonstrated that FtaC is responsible for the methylation of CDP-Thr, using SAM as the methyl donor. We believe that the methylation event is not the rate limiting step in the Type A glycan biosynthesis as we only found a very small amount of CDP-Thr in an *ftaD* mutant strain, while it accumulated in a *ftaC* mutant strain. The identification of FtaC as a CDP-threonine methyltransferase extends the large family of SAM-dependent methyltransferases (EC.2.1.1.X, currently more than 400 members). Based on the AlphaFold (v 2.0)-predicted structure (54), FtaC belongs to the class I methyltransferases, which typically contain a seven-stranded β-sheet flanked by α-helices (60-62). In fact, it appears to only contain the consensus structural core, without any auxiliary domains and therefore represents one of the smallest methyltransferases described to date.

We could couple the FtaC and FtaD activity in a single reaction, mirroring the last two steps of the Type A biosynthesis in vitro. Unfortunately, so far we have been unable to purify FtaB, the key enzyme for CDP-Thr synthesis because it repeatedly ended up in the insoluble fraction. Interestingly, AlphaFold (v 3.0) (54) predicts heterodimerization of FtaA and FtaB, indicating that these enzymes form a complex in *C. difficile*, possibly explaining the FtaB solubility problems during purification from *E. coli*. In the future, by incorporating purified FtaA and FtaB, we aim to recapitulate the full PGM biosynthesis.

In *P. aeruginosa* PAO1, a glycan structure (b-type) similar to the *C. difficile* Type A glycan is found (45). This structure also contains a monosaccharide (most probably a rhamnose (63)) which is linked to a *N,N-*dimethylated threonine (32) through a phosphodiester bond. Interestingly, in the methyltransferase mutant in *P. aeruginosa*, no PGM was observed, and FliC was only decorated with the deoxyhexoses, suggesting that methylation is required for the phosphotransferase. Conversely, methylation is not an absolute requirement for FtaD activity although the data from our in vitro experiments clearly showed that *C. difficile* FtaD strongly prefers CDP-Me-Thr over CDP-Thr as the donor substrate.

Given the dimethylation of the threonine in the *P. aeruginosa* b-type glycan, we predicted that CDP-*N,N*-dimethylthreonine (CDP-diMe-Thr) is the endogenous substrate of the phosphotransferase PA1091 in *P. aeruginosa* (32). Previously, we showed a minor fraction of the *C. difficile* Type A glycan with a dimethylated threonine (44). Notably, our biochemical experiments with purified FtaC provided no evidence for the formation of CDP-diMe-Thr under these conditions. These data imply that in *C. difficile* an additional, FtaC-independent methylation of CDP-Me-Thr might exist or that dimethylation *in vivo* is dependent on conditions not recapitulated in our *in vitro* assay.

We have also identified biosynthetic gene clusters that encode part of the FtaABCD cluster in other organisms, of which those that combine FtaB and FtaD are likely capable of synthesizing CDP-threonine as an intermediate to make a structure that includes phosphothreonine. It would be interesting to see if CDP-threonine can be detected in these species, perhaps serving as an intermediate in the synthesis of other biomolecules. In our analysis, FtaB homologues were mostly associated with a flagellar locus, suggesting a role in flagellin glycan modifications. However, we have also found a FtaB homolog in the non-flagellated *Methanobrevibacter* (Archae). This protein is encoded in a region that seems to be involved in lipid biosynthesis, possibly implicating that CDP-threonine might also be involved in modification of glycolipids.

In conclusion, we have unravelled the biosynthetic pathway involved in the Type A glycan modification of the important virulence factor FliC in the human opportunistic pathogen *C. difficile* with implications for post-glycosylation modifications in other Bacillota and Pseudomonadota. Most importantly, we describe the role of novel cellular CDP-amino acid conjugates, CDP-threonine and CDP-*N*-methylthreonine. Their identification may imply that such high-energy intermediates may play a more general role in cellular pathways than previously anticipated.

## Materials and Methods

### Materials

Synthetic peptides with a Ser-β-D-GlcNAc, that were used as acceptor substrate peptides for FtaD, were synthesized in-house using standard Fmoc solid-phase peptide synthesis. For the details of the synthesis and all characterization data of the synthetic intermediates CDP-threonine and CDP-*N-* methylthreonine, see Supporting Information.

### Plasmids

Plasmids were ordered from Twist biosciences. They consisted of a pET28b backbone with the codon-optimized sequences of *ftaC* (pJC120) and *ftaD* (pJC121), both with N-terminal His6 tags cloned between NdeI and XhoI. These plasmids were used for overexpression in *E. coli*.

### Bacterial culture and media

*C. difficile* strains were cultured in brain-heart infusion (BHI, Oxoid) broth supplemented with 0,5 % yeast extract (BHIY, Sigma), or on BHIY plates (1.5 % agar (Thermo)) at 37 °C in an anaerobic Whitley A55 HEPA Workstation (Don Whitley Scientific, Yorkshire, UK). *E. coli* strains were grown in Lysogeny Broth (LB) supplemented with appropriate antibiotics. *C. difficile* strains used in this study are shown in Table 1.

**Table 1:** C. *difficile* strains used in this study.

| Name | Description | Genotype | Flagellin<br>modification Type | Reference |
| --- | --- | --- | --- | --- |
| WKS2044 | 630Δ*erm* | 630Δ*erm* (PCR ribotype 012) | A | (29, 31) |
| WKS2047 | *ftaA* mutant | 630Δ*erm*, *cd0241::CT* | A | (29, 31) |
| WKS2049 | *ftaB* mutant | 630Δ*erm*, *cd0242::CT* | A | (29, 31) |
| WKS2051 | *ftaC* mutant | 630Δ*erm*, *cd0243::CT* | A | (29, 31) |
| WKS2052 | *ftaD* mutant | 630Δ*erm*, *cd0244::CT* | A | (29, 31) |
| WKS2045 | *fliC* mutant | 630Δ*erm*, *fliC::CT* | A | (29) |
| WKS2036 | *fgtA* mutant | 630Δ*erm*, *cd0240::CT* | A | (29) |
| LL-RT017 | LL-RT017 | PCR ribotype 017 | A | (41, 68) |
| LL-RT027 | R20291 | PCR Ribotype 027 | B | (41, 68) |

### Recombinant protein expression and purification

Expression of N-terminally His6-tagged FtaC was induced with 1 mM IPTG in *E. coli* Rosetta cells in mid-exponential growth phase (optical density approximately 0.5). After 3-4 h of induction, soluble recombinant protein was purified using standard techniques. Briefly, cells were pelleted using centrifugation and resuspended in lysis buffer (50 mM NaH_2_PO_4_, 300 mM NaCl, 10 mM Imidazole, pH

8). Subsequently, cells were lysed by sonication and cleared lysate was incubated with 1 mL Ni-sepharose (GE healthcare). After 2 hr incubation while mixing, the Sepharose beads were washed with 20 mM and 50 mM imidazole in lysis buffer and finally the protein was eluted using 250 mM imidazole in lysis buffer.

Expression of N-terminally His6-tagged FtaD was induced with 1 mM IPTG in *E. coli* Rosetta cells in mid-exponential growth phase (OD=0.6). After overnight induction at 30°C, cells were pelleted using centrifugation, cell pellet was resuspended in lysis buffer (50 mM TRIS-Cl, 300 mM NaCl, 10 mM imidazole, pH=8.0, cOmplete^™^ Protease Inhibitor Cocktail without EDTA, lysozyme) and lysed by sonication. To the lysate DNAse 1 was added to 5 µg/mL final concentration, lysate was incubated on ice for 15 minutes and cleared by centrifugation. Supernatant was purified over a 5mL Cytiva HisTrap column (50mM TRIS-Cl, 300 mM NaCl, 10mM to 160 mM imidazole, pH=8.0) and fractions containing FtaD were concentrated over a 15 mL Amicon 10 kDa cutoff spin filter. Finally, buffer was exchanged (20 mM sodium-HEPES, 200 mM NaCl, pH=7.6) and protein was purified using size exclusion chromatography (Superdex 200 10/300 GL, protein at about 0.5 CV, 1 CV=20mL). Fractions containing protein were combined, concentrated over a 15 mL Amicon 10 kDa cutoff spin filter and diluted to a final concentration of 1 mg/mL, snap-frozen in liquid nitrogen, and stored at -80°C in 30 µL aliquots until further use.

### *C. difficile* metabolite extraction

*C. difficile* cells from 25 mL cultures at an OD_600nm_ of 0.8 were pelleted and resuspended in 25 mL PBS and divided in 2 mL aliquots. Cell pellets from these aliquots were snap-frozen in liquid nitrogen and stored at -80 °C until further use.

For metabolite extraction, pellets were reconstituted in 0.5 mL of ice-cold LC-grade MeOH (LiChrosolv®, Supelco)/milliQ H_2_O (80:20 (v/v)). Subsequently, samples were sonicated (three times 30sec and placed on ice in between steps) and centrifuged (14000 × g, 5 min, 4 °C). Supernatants were collected and the pellets were used for a second round of extraction as described above, and supernatants were combined and freeze-dried. Then, 945µL LC-grade MeOH, 105µL chloroform (Emsure®, Supelco), and 350µL LC-grade H_2_O (CHROMASOLV LC-MS Ultra, Honeywell) was added to each freeze-dried sample, vortexed for 20 seconds and placed onto dry ice for 20 seconds. This step was repeated 3 times. Finally, the samples were centrifuged (14000 × g, 15 min, 0 °C) and supernatants were collected and dried under nitrogen gas. Samples were dissolved in 100 µL LC-grade ACN (CHROMASOLV™, Honeywell)/LC-grade H_2_O (50:50 (v/v)).

### HILIC-MS/MS

Hydrophilic interaction chromatography-tandem mass spectrometry (HILIC-MS/MS) was performed using a LC-30 chromatography system (Shimadzu) coupled to a TripleTOF 6600+ mass spectrometer (Sciex). Aliquots of 2 µL from each sample (in ACN/H_2_O, 1:1 (v/v)) were co-injected with 8µL of ACN on a iHILIC-(P) Classic column (100 x 2.1 mm, 5 µm, 200 Å). The column oven temperature was set at 30°C. Mobile phases consisted of solvent A (20 mM ammonium acetate + 20 mM ammonium hydroxide in LC-grade water) and B (100% ACN). Metabolites were eluted using the following gradient: 0 -1.5 min: 90% B, 1.5-8 min: 90-50% B, 8-11 min: 50% B, 11-12 min: 50-45% B, 12-15 min: 45% B, 15-15.01 min: 45-90% B, 15.01-24 min: 90% B. The flow rate was 200 µL/min. The mass spectrometer was operated in negative ion mode (spray voltage -4500 V) with a scan range of *m/z* 50-900. MS/MS was performed on predefined precursor ions corresponding to CPD-threonine (CDP-Thr, *m/z* 503.059, [M-H]^-^) and CDP-*N*-methylthreonine (CDP-Me-Thr, *m/z* 517.074, [M-H]^-^). Stock solutions of CDP-Thr and CDP-Me-Thr were prepared at 1-2 µg/mL.

### Proteomics analysis

Protein extracts were generated and digested with trypsin as described previously (31). Tryptic peptides were dissolved in water/formic acid (100:0.1 (v/v)) and analyzed by on-line C18 nanoHPLC MS/MS with a system consisting of an Ultimate3000nano gradient HPLC system (Thermo, Bremen, Germany), and an Exploris480 mass spectrometer (Thermo). Fractions were injected onto a cartridge precolumn (300 μm × 5 mm, C18 PepMap, 5 um, 100 A, and eluted via a homemade analytical nano-HPLC column (30 cm × 75 μm; Reprosil-Pur C18-AQ 1.9 um, 120 A (Dr. Maisch, Ammerbuch, Germany). The gradient was run from 2% to 36% solvent B (H_2_O/ACN/formic acid (20:80:0.1 (v/v/v)) in 30 min at 250 nl/min. The nano-HPLC column was drawn to a tip of ∼10 μm and acted as the electrospray needle of the MS source. The mass spectrometer was operated in a data-dependent cycle time mode of 3 seconds, with a normalized HCD collision energy of 30% and recording of the MS2 spectrum in the orbitrap, with a quadrupole isolation width of 1.2 Th. In the master scan (MS1) the resolution was 120,000, the scan range *m/z* 350-1600, at standard AGC target and a maximum fill time of 50 ms. A lock mass correction on the background ion at *m/z*=445.12003 was used. Precursors were dynamically excluded after n=1 with an exclusion duration of 10 s, and tolerance of 10 ppm. Included charge states were 1-5. For MS2 the first mass was set to 110 Da, and the MS2 scan resolution was 30,000 at an AGC target of 100% at a maximum fill time of 60 ms.

### Biochemical assays

Typical FtaC methyltransferase assays were performed for 60 min at 37°C in a buffer containing 25 mM Tris (pH 7.5), 10 mM EDTA, 1 mM dithiothreitol (DTT), 200 µM *S*-adenosylhomocysteine (SAM), 100 µM CDP-threonine and 1.4 µg of purified His6-tagged FtaC. Subsequently, samples were analyzed using the MTase-Glo™ Methyltransferase Assay (Promega, Madison, Wisconsin), following the manufacturer’s instruction. Bioluminescence intensity was measured using an EnVision 2105 Multi-Mode Microplate Reader (Perkin Elmer, Waltham, Massachusetts) or by HILIC-MS, as described above.

FtaD assays were performed in 100 mM Tris-HCl buffer (pH 7.6) containing 200 µM CDP-(Me)-threonine, 40 µM acceptor substrate peptide (peptides with a Ser-β-D-GlcNAc) and 2 µg recombinant His6-tagged FtaD, at 37°C. After incubation, peptides were recovered by reversed-phase solid-phase extraction cartridges (Oasis HLB 1⍰cm^3^ 10⍰mg; Waters, Etten-Leur, the Netherlands) and eluted with 300]μL of ACN/0.1% FA (35:65 (v/v). Samples were diluted till a peptide concentration of 100 fmol/µL and subsequently analyzed by online C18 nanoHPLC MS/MS with a system consisting of an Easy nLC 1200 (Thermo, Bremen, Germany) coupled to a LUMOS mass spectrometer (Thermo) or a system consisting of an Ultimate 3000 (Thermo) coupled to an Exploris mass spectrometer (Thermo). Fractions were injected onto a home-made precolumn (100 μm × 15 mm; Reprosil-Pur C18-AQ 3 μm, Dr. Maisch, Ammerbuch, Germany) and eluted via a home-made analytical nano-HPLC column (30 cm × 75 μm; Reprosil-Pur C18-AQ 1.9 um). The gradient was run from 2% to 40% solvent B (H_2_O/acetonitrile/formic acid (20:80:0.1 (v/v/v)) in 30 min. The nano-HPLC column was drawn to a tip of ∼5 μm and acted as the electrospray needle of the MS source. The mass spectrometer was operated in data-dependent MS/MS mode for a cycle time of 3 seconds, with an HCD collision energy at 32 V and recording of the MS2 spectrum in the orbitrap. In the master scan (MS1) the resolution was 120,000, the scan range *m/z* 400-1500, at an AGC target of 400,000 @maximum fill time of 50 ms. Dynamic exclusion after n=1 with exclusion duration of 10 s. Charge states 2-5 were included. For MS2 precursors were isolated with the quadrupole with an isolation width of 1.2 Th. First mass was set to m/z 110. The MS2 scan resolution was 30,000 with an AGC target of 50,000 at maximum fill time of 60 ms.

### Analysis of gene clusters containing ftaA-ftaD homologs

To search for gene clusters of *fliC, fgtA* and *ftaA-ftaD* (*cd0241-cd0244)*, bacterial genomes were downloaded from the mOTUs 3.1.1. reference database (64). Gene calling was run using Pyrodigal v3.5.1, (65, 66) in meta mode (-p meta) to obtain all protein sequences. Subsequently, a database was prepared with cblaster makedb v1.3.20 (67). To detect the clusters, a cblaster search was run with the protein sequences of FliC, FgtA and FtaA-FtaD, requesting two unique hits, a hitlist size of 900M and DIAMOND in sensitive mode. The hits were combined with the mOTUs 3.1.0 metadata (https://zenodo.org/records/10275750) into a single table (Supplementary Table 1).

## Acknowledgements

This research was funded by the research program Investment Grant NWO Medium with project number 91116004, which is (partially) financed by ZonMW. We thank dr. Brendan Wren (London School of Hygiene and Tropical Medicine) for providing the *C. difficile* mutant strains. We thank dr. Jan Wouter Drijfhout for helpful discussions. Z.A. acknowledges the Dutch Research Council (NWO) for funding support (VI.Veni.212.173)

