## Supporting information for "Identification of novel cellular intermediates unveils unique enzymes for flagellar glycan biosynthesis in *Clostridioides difficile*"

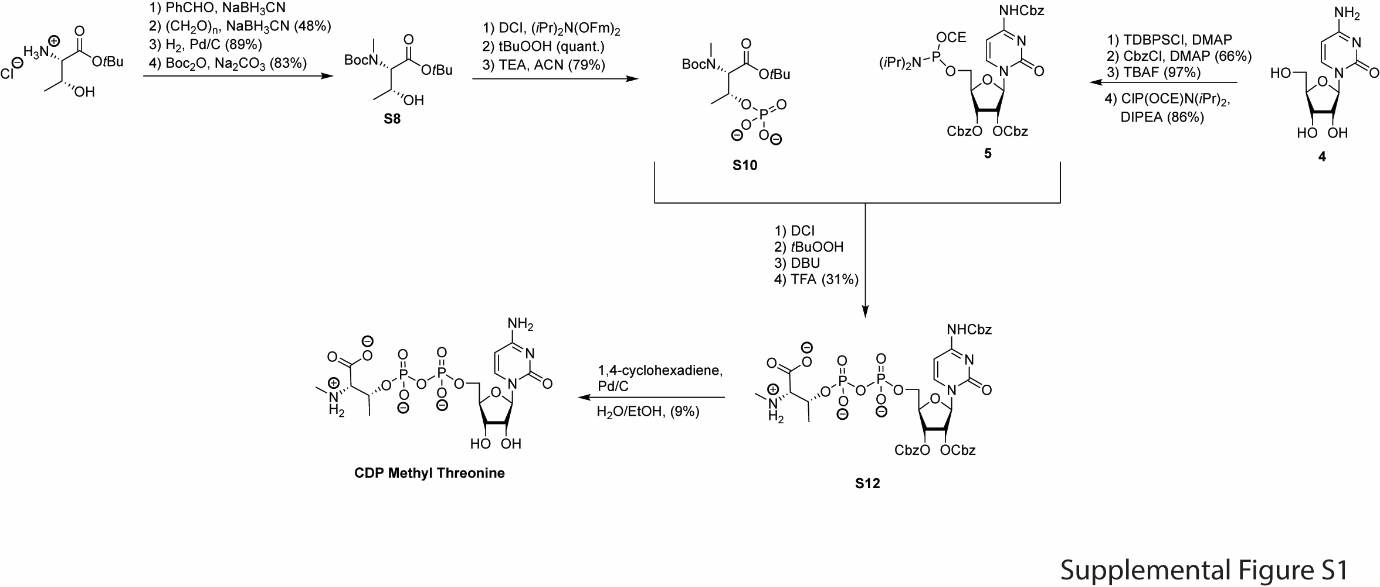

**Supplemental Figure S1: Synthesis and characterization of CDP-*N*-methylthreonine (CDP-Me-Thr).**

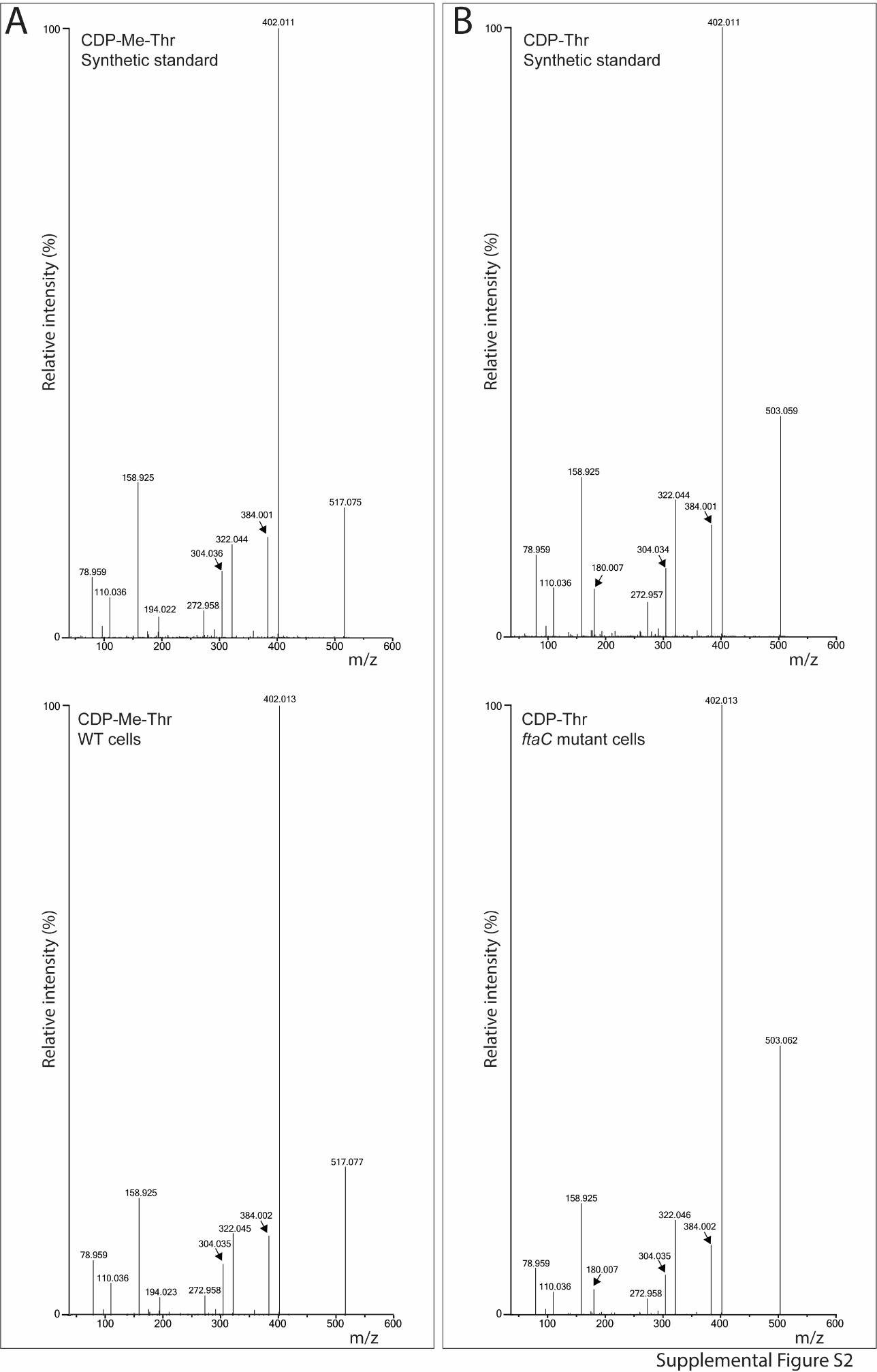

**Supplemental Figure S2: Comparison of MS/MS spectra of synthetic and endogenous CDP-*N*-methylthreonine (CDP-Me-Thr, A) and CDP-threonine (CDP-Thr, B).**

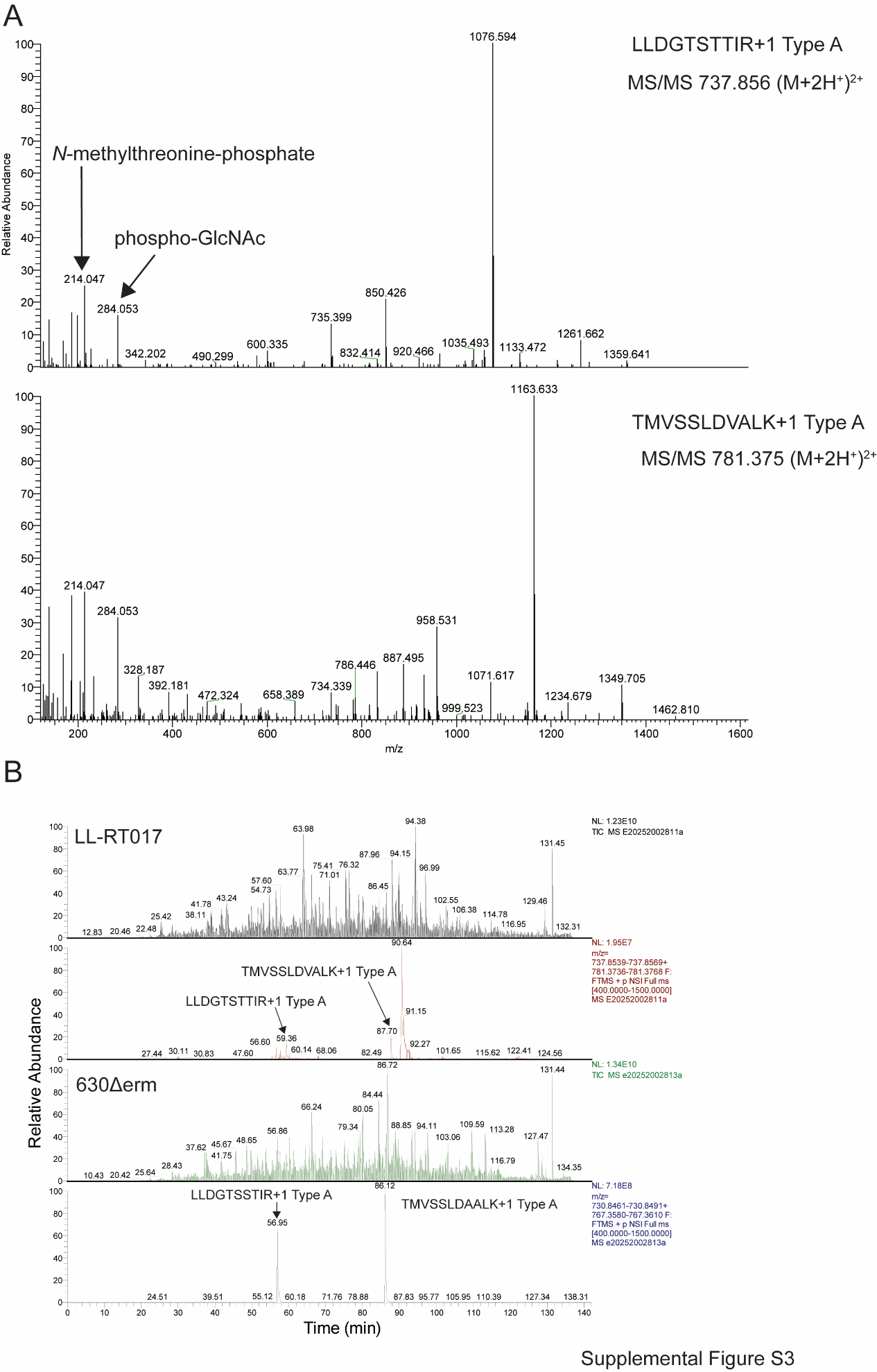
**Supplemental Figure S3: Analysis of Type A-modified peptides in a *C. difficile* strain LL-RT017 and 630Δ*erm*.** A: MS/MS spectra of Type A-modified peptides in strain LL-RT017. B: Total ion chromatogram and extracted ion chromatograms of ions corresponding to Type A-modified peptides in strain LL-RT-017 (upper panels) and 630Δ*erm* (lower panels).

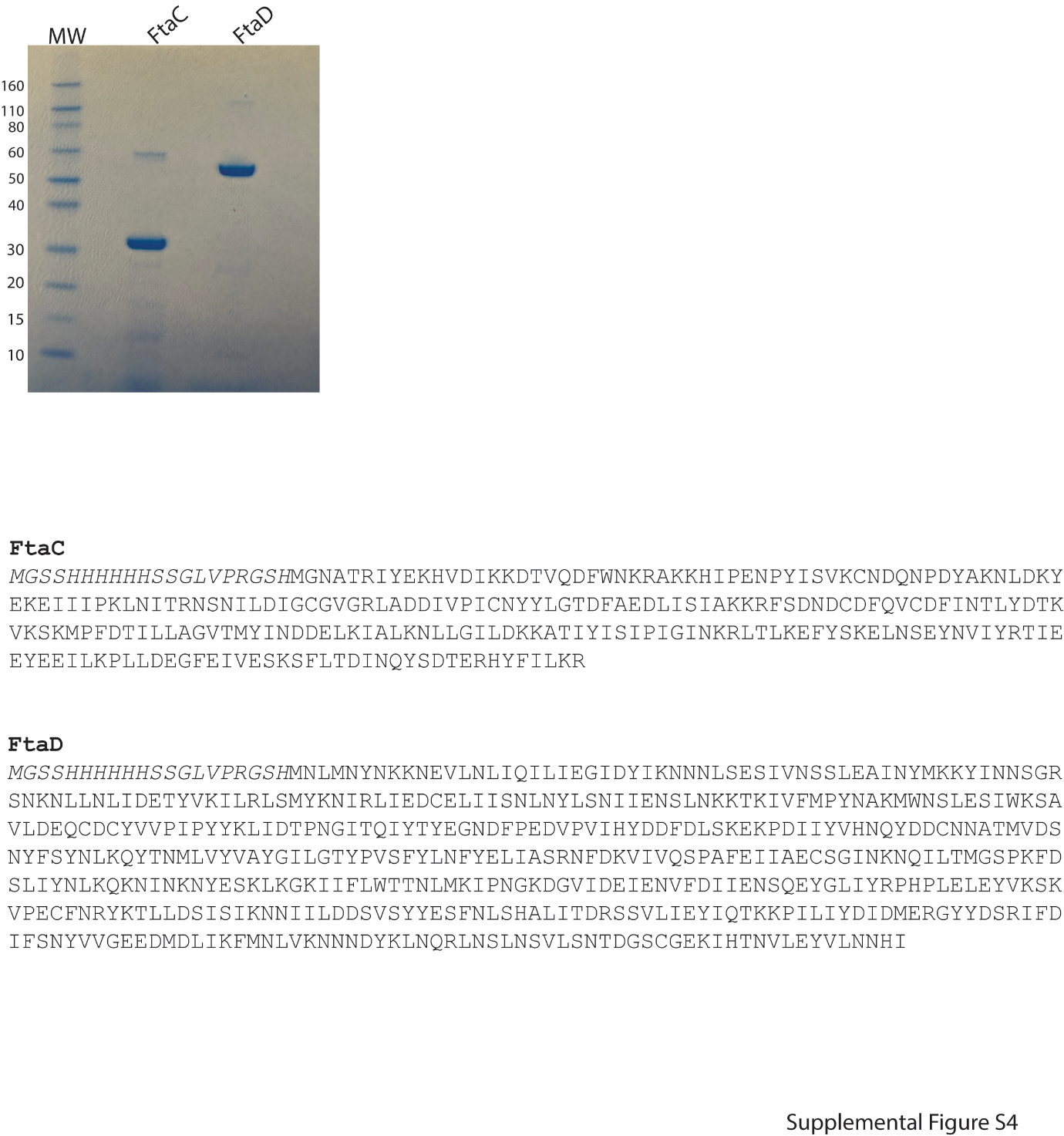
**Supplemental Figure S4: SDS-PAGE analysis and sequence of recombinant FtaC and FtaD**

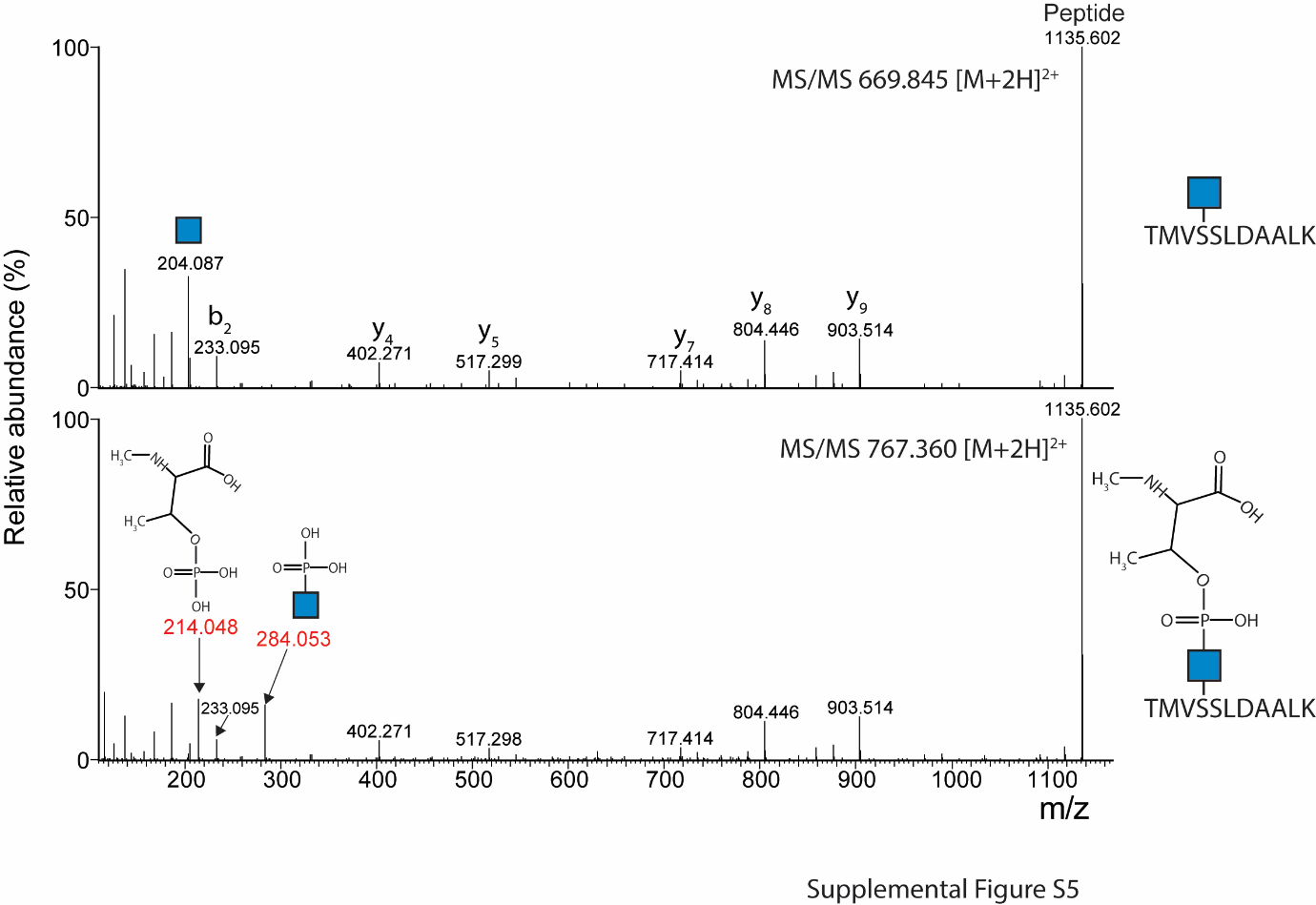
**Supplemental Figure S5: MS/MS spectra of the substrate and product peptide of a FtaD assay.**

FtaD was incubated with the donor substrate CDP-Me-Thr and the substrate acceptor peptide TMVsSLDAALK (s=Ser-β-D-GlcNAc). LC-MS/MS was used to characterize the substrate acceptor peptide (upper panel) and product peptide (lower panel), i.e., the peptide with the full Type A structure.

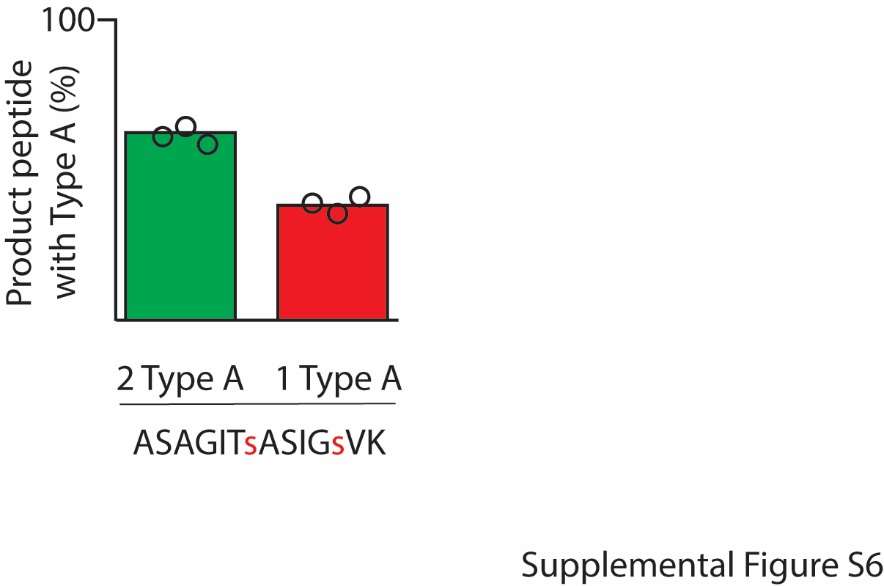

**Supplemental Figure S6: FtaD activity assay with a synthetic peptide with two *O*-GlcNAcs.**

FtaD was incubated with the donor substrate CDP-Me-Thr and the acceptor substrate peptide ASAGITsASIGsVK (s=Ser-β-D-GlcNAc). LC-MS/MS was then performed to determine the relative abundances of the two product peptides, i.e., the peptide where a single or both serines were modified with a Type A structure.

**Supplemental data chemistry**

**Cytidine and threonine building block**

**General experimental**

All chemicals were used as received unless stated otherwise. Molecular sieves were activated by flame drying *in vacuo* before use. Solvents were dried over activated 4 Å molecular sieves except for MeCN, which was dried over 3 Å molecular sieves. Reactions were performed under a N_2_ atmosphere unless stated otherwise. Reaction mixtures were concentrated *in vacuo* using rotary evaporators at 40-50 °C unless stated otherwise. Reactions were monitored by thin layer chromatography (TLC) analysis using silica gel 60 F254 coated aluminium sheets from Merck. TLC plates were visualized with ultraviolet light (254 nm) or sprayed with H_2_SO_4_ (20% v/v in MeOH), potassium permanganate (1 g KMnO_4_, 5 g K_2_CO_3_, in 200 ml H_2_O) or ceric ammonium molybdate (1 g Ce(NH_4_)_4_(SO_4_)_4_•2H_2_O, 2.5 g (NH_4_)_6_Mo_7_O_24_•4H_2_O, 10 ml H_2_SO_4_ in 90 ml H_2_O). ^13^C NMR spectra are acquired via the attached proton test (APT) experiment and are presented with even signals (C_q_ and CH_2_) pointing upwards and odd signals (CH and CH_3_) pointing downwards. ^13^C NMR spectra are proton decoupled. The chemical shifts are noted as δ-values in parts per million (ppm) relative to the tetramethylsilane signal (TMS, δ = 0 ppm) or the corresponding solvent signal for ^1^H NMR and relative to the corresponding solvent signal for ^13^C NMR. Phosphorylation reactions were monitored with ^31^P NMR using an acetone-D_6_ insert for a locking signal, and the resulting spectra were indirectly calibrated with H_3_PO_4_. Structural assignments were made with additional information from gCOSY, gHSQC, and gHMBC experiments. HRMS samples were prepared in either MeCN or milliQ grade H_2_O and diluted in a solution of MeCN/milliQ grade H_2_O/*t*BuOH (1/1/1) with an approximate concentration of 1 mM and measured on a Thermo Scientific LTQ Orbitrap XL.

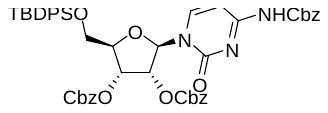
***N*^4^-benzyloxycarbonyl-2’,3’-di-*O*-(benzyloxycarbonyl)-5’-*O*-t*ert*-butyldiphenylsilyl cytidine (S1)**

Cytidine (4.9 g, 20 mmol, 1 eq.) and DMAP (0.49 g, 4 mmol, 0.2 eq.) were combined, co-evaporated thrice with anhydrous pyridine, and dissolved in anhydrous pyridine (100 mL, 0.2 M). The solution was cooled to 0°C, and TBDPS-Cl (5.4 mL, 21 mmol, 1.05 eq.) was dropped to the stirring solution. The reaction mixture was stirred overnight at rt until TLC analysis (DCM:MeOH 7:3) indicated full conversion. Upon conversion, the reaction mixture was diluted in DCM, washed with NaHCO_3_ (sat.aq.) and the aqueous layer was extracted with DCM (3x). The combined organic layers were washed with brine (1x), dried over Na_2_SO_4_, filtered and concentrated *in vacuo*. The resulting residue was taken up in anhydrous DCM (200 mL, 0.1 M), DMAP (11.0 g, 90 mmol, 4.5 eq.) was added, the solution was cooled to 0°C and benzyl chloroformate (11.4 mL, 80 mmol, 4 eq.) was dropped to the mixture. The reaction mixture was stirred overnight at rt. When TLC analysis (pentane:EtOAc 6:4) indicated full conversion, the reaction was quenched with H_2_O and the organic layer was separated. The aqueous layer was back extracted with EtOAc (3x), the combined organic layers were washed with brine (1x), dried over Na_2_SO_4_ and concentrated *in vacuo*. Column chromatography (pentane:EtOAc 6:4) yielded the desired product as a white solid over 2 steps (11.71 g, 13.3 mmol, 66%). **^1^H NMR** (400 MHz, CDCl_3_) δ 8.23 (d, *J* = 7.6 Hz, 1H), 7.70 – 7.57 (m, 4H), 7.47 – 7.30 (m, 21H), 7.01 (d, *J* = 7.5 Hz, 1H), 6.25 (d, *J* = 3.1 Hz, 1H), 5.48 – 5.39 (m, 2H), 5.21 (s, 2H), 5.15 (s, 2H), 5.11 – 5.05 (m, 2H), 4.30 (d, *J* = 6.0 Hz, 1H), 4.17 – 3.75 (m, 2H), 1.11 (s, 9H). **^13^C NMR** (101 MHz, CDCl_3_) δ 162.5, 154.8, 153.9, 143.9, 135.7, 135.5, 131.9, 128.9, 128.2, 95.3, 88.0, 81.5, 77.5, 77.2, 72.2, 70.5, 68.1, 61.9, 27.1, 19.4. **HRMS** (ESI) m/z: [M + H]^+^ Calcd for C_49_H_50_N_3_O_11_Si^+^ 884.3209; Found 884.3202.

***N*^4^-benzyloxycarbonyl-2’,3’-di-*O*-(benzyloxycarbonyl) cytidine**
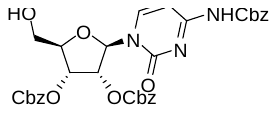
 **(S2)**

Compound **S1** (11.71 g, 13.3 mmol, 1 eq.) was dissolved in THF (133 mL, 0.1 M) and cooled to 0°C after which AcOH (1.14 mL, 20 mmol, 1.5 eq.) and TBAF (1 M in THF; 20 mL, 20 mmol, 1.5 eq.) were added. The reaction mixture was allowed to stir overnight at rt until TLC analysis revealed full conversion. Upon completion, H_2_O was added to the solution and the aqueous layer was extracted with EtOAc (3x), the organic layer was washed with brine (1x), dried over Na_2_SO_4_, filtered and concentrated *in vacuo*. Column chromatography (DCM/MeOH, 100/0🡪98/2) furnished the titled compound (8.29 g, 12.90 mmol, 97%) as a white solid. **^1^H NMR** (400 MHz, CDCl_3_) δ 7.88 (d, *J* = 7.5 Hz, 1H), 7.74 (s, 1H), 7.37 (s, 5H), 7.34 (s, 5H), 7.32 (s, 5H), 7.28 (d, *J* = 7.5 Hz, 1H), 5.81 – 5.74 (m, 2H), 5.54 – 5.47 (m, 1H), 5.20 (s, 2H), 5.12 – 5.08 (m, 4H), 4.33 (dt, *J* = 4.3, 2.0 Hz, 1H), 4.02 – 3.95 (m, 1H), 3.86 – 3.74 (m, 1H), 3.71 – 3.61 (m, 1H).**^13^C NMR** (101 MHz, CDCl_3_) δ 163.0, 154.2, 153.8, 152.2, 146.8, 134.7, 128.9, 128.8, 128.8, 128.7, 128.6, 128.6, 128.6, 95.8, 92.8, 83.8, 75.7, 73.8, 70.6, 70.5, 68.3, 61.5. **HRMS** (ESI) m/z: [M + H]^+^ Calcd for C_33_H_32_N3O_11_^+^ 646.2031; Found 646.1999.

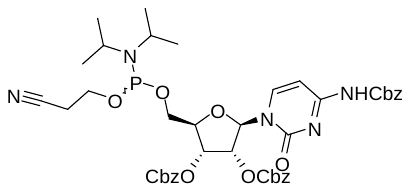
***N^4^*-benzyloxycarbonyl-2',3'-bis-*O*-(benzyloxycarbonyl)cytidine 5'-O-(β-cyanoethyl *N,N*-diisopropylphosphoramidite) (5)**

Compound **S3** (324 mg, 0.50 mmol, 1 eq.) was co-evaporated with toluene (3x) before being dissolved in dry DCM (5 mL, 0.1 M). DiPEA (0.26 mL, 1.50 mmol, 3 eq.) and 2-cyanoethyl *N,N*-diisopropylchlorophosphoramidite (0.15 mL, 0.55 mmol, 1.1 eq.) were subsequently added and the reaction mixture was stirred at rt for 2 h until TLC analysis revealed full conversion. The reaction mixture was then diluted in DCM, washed with brine (1x), dried over Na_2_SO_4_, filtered and concentrated *in vacuo*. Column chromatography (toluene/acetone/TEA, 100/0/1🡪95/5/1) yielded phosphoramidite **5** (365 mg, 0.43 mmol, 86%) as *R*p and *S*p diastereomeric mixture and as a white foamy solid. **^1^H NMR** (400 MHz, CD_3_CN) δ 8.53 (s, 1H), 8.09 (2x d, *J* = 7.7 Hz, 1H), 7.47 – 7.31 (m, 10H), 7.28 – 7.22 (m, 3H), 7.21 – 7.12 (m, 2H), 6.05 (2x d, *J* = 4.3 Hz, 1H), 5.43 – 5.30 (m, 2H), 5.20 (s, 2H), 5.19 – 5.03 (m, 4H), 4.43 – 4.35 (m, 1H), 4.02 – 3.73 (m, 4H), 3.70 – 3.53 (m, 2H), 2.70 – 2.59 (m, 2H), 1.25 – 1.11 (m, 12H). **^13^C NMR** (101 MHz, CD_3_CN) δ 164.1, 154.8, 154.7, 145.6, 136.8, 136.2, 136.1, 129.9, 129.6, 129.6, 129.3, 129.3, 129.2, 129.0, 126.2, 95.9, 89.6, 89.1, 82.8, 82.7, 82.5, 82.4, 78.1, 74.8, 74.7, 71.1, 71.0, 68.3, 63.4, 63.2, 63.1, 59.9, 59.7, 59.7, 59.5, 44.1, 44.0, 43.9, 43.9, 25.0, 25.0, 24.9, 21.4, 21.0, 21.0, 20.9. **^31^P NMR** (162 MHz, CD_3_CN) δ 149.1, 148.8. **HRMS** (ESI) m/z: [M + H]^+^ Calcd for C_36_H_36_N_4_O_13_P^+^ (H-phosphonate) 763.2011, Found 763.2008.

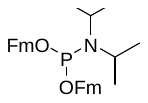
***N*,*N*-diisopropyl bis(9-methylfluorenyl)phosphoramidite (S4)**

Compound **S4** was prepared according to literature.^1^ PCl_3_ (1.00 mL, 11.40 mmol, 1 eq.) was added to dry THF (40 mL, 0.28 M) in a flame dried flask. The mixture was cooled to 0°C before adding DiPEA (4.00 mL, 23.00 mmol, 2 eq.) and DIPA (with a syringe pump over 10 minutes; 3.04 mL, 21.66 mmol, 1.9 eq.). The reaction mixture was stirred for 1 h at 0°C before adding another portion of DiPEA (4.00 mL, 23.00 mmol. 2 eq.) and FmOH (2.3 M in dry THF; 10 mL, 22.80 mmol, 2 eq.) at 0°C. The reaction mixture was allowed to stir overnight at rt before concentrating *in vacuo*. The crude was diluted in EtOAc and washed with phosphate buffer solution pH 7 (50 mL). The aqueous layer was extracted with EtOAc (4x), washed with phosphate buffer solution pH 7 (50 mL), dried over Na_2_SO_4_, filtered and concentrated *in vacuo*. Column chromatography (pentane/EtOAc/TEA, 99/1/1🡪98/2/1) afforded phosphoramidite **S4** (4.49 g, 8.61 mmol, 76%) as a colourless oil. **^1^H NMR** (300 MHz, CD_3_CN) δ 7.81 – 7.71 (m, 4H), 7.55 (d, *J* = 7.5 Hz, 4H), 7.42 – 7.20 (m, 8H), 4.04 (t, *J* = 6.1 Hz, 2H), 3.95 – 3.81 (m, 2H), 3.78 – 3.63 (m, 2H), 3.58 – 3.35 (m, 2H), 1.05 (d, *J* = 6.8 Hz, 12H). **^31^P NMR** (121 MHz, CD_3_CN) δ 148.3.

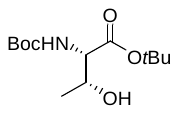
***N*-*tert*-butyloxycarbonyl-l-threonine *tert*-butyl ester** **(2)**

Boc-Thr-OH (1.75 g, 8.00 mmol, 1 eq.) was dissolved in dry DCM (80 mL, 0.1 M) and *tert*-butyl *N*,*N'*-diisopropylcarbamimidate (7.2 mL, 32.00 mmol, 4 eq.) was added. The reaction mixture was stirred overnight at rt until TLC analysis revealed full conversion (upon completion a white precipitate form). The mixture was filtered over Celite, the residue was washed with DCM and concentrated *in vacuo*. Column chromatography (pentane/EtOAc, 95/5🡪80/20) provided the titled compound (2.03 g, 7.37 mmol, 92%) as a colourless oil. **^1^H NMR** (400 MHz, CDCl_3_) δ 5.28 (d, *J* = 8.9 Hz, 1H), 4.28 – 4.17 (m, 1H), 4.12 (d, *J* = 7.9 Hz, 1H), 2.17 (d, *J* = 5.7 Hz, 1H), 1.47 (s, 9H), 1.44 (s, 9H), 1.22 (d, *J* = 6.4 Hz, 3H). **^13^C NMR** (101 MHz, CDCl_3_) δ 170.7, 156.3, 82.5, 80.0, 68.6, 59.3, 28.4, 28.1, 20.1.

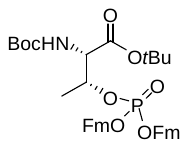
***N*-(*tert*-butoxycarbonyl)-*O*-(bis((9H-fluoren-9-yl)methoxy))phosphoryl-l-threonine *tert*-butyl ester (S5)**

Threonine **2** (550 mg, 2.00 mmol, 1 eq.) and phosphoramidite **S4** (1.15 g, 2.20 mmol, 1.2 eq.) were co-evaporated with toluene (3x) before being dissolved in dry MeCN (24 ml, final total concentration is 0.1 M). Freshly activated 3Å molecular sieves were added and the mixture was allowed to stir 30 minutes at rt before adding DCI (0.25 M in MeCN; 16 mL, 4.00 mmol, 2 eq.). The reaction mixture was stirred for another 30 minutes until ^31^P NMR revealed full conversion. Upon completion, *t*BuOOH (5-6 M in decane, 1.09 mL, 6.00 mmol, 3 eq.) was added and the reaction mixture was stirred for 1 h until ^31^P NMR revealed full oxidation. NaHCO_3_ (sat. aq.) was added and the aqueous layer was extracted with EtOAc (3x), washed with brine (1x), dried over Na_2_SO_4_, filtered and concentrated *in vacuo*. Column chromatography (pentane/EtOAc, 85/15🡪75/25) provided phosphotriester **S5** (1.32 mg, 1.89 mmol, 94%) as a foamy solid. **^1^H NMR** (400 MHz, CDCl_3_) δ 7.76 – 7.66 (m, 4H), 7.57 – 7.46 (m, 3H), 7.44 – 7.28 (m, 7H), 7.24 – 7.22 (m, 1H), 7.19 – 7.13 (m, 1H), 5.15 (d, *J* = 9.5 Hz, 1H), 4.92 – 4.81 (m, 1H), 4.33 – 4.24 (m, 1H), 4.18 (m, 4H), 4.13 – 4.04 (m, 2H), 1.45 (s, 9H), 1.42 (s, 9H), 1.23 (d, *J* = 6.4 Hz, 3H). **^13^C NMR** (101 MHz, CDCl_3_) δ 168.9, 143.1, 141.5, 129.2, 128.0, 128.0, 127.2, 125.3, 125.2, 125.2, 120.1, 120.1, 82.9, 80.8, 76.1, 69.3, 69.3, 69.2, 58.5, 48.0, 28.4, 28.1, 18.5. **^31^P NMR** (162 MHz, CDCl_3_) δ -2.7. **HRMS** (ESI) m/z: [M + Na]^+^ Calcd for C_41_H_46_NO_8_PNa^+^ 734.2853; Found 734.2851.

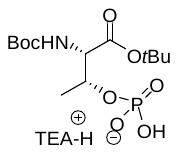
***N*-(tert-butoxycarbonyl)-*O*-phosphono-l-threonine *tert*-butyl ester triethylammonium salt (3)**

Phosphotriester **S5** (1.32 g, 1.89 mmol, 1 eq.) was dissolved in MeCN (18.9 mL, 0.1 M) before adding TEA (3.95 mL, 28.35 mmol, 15 eq.). The reaction mixture was stirred overnight at rt until ^31^P NMR revealed full conversion to the monophosphate. The mixture was then concentrated *in vacuo* and column chromatography (DCM/MeOH, 98/2🡪85/15) furnished phosphate **3** (0.68 g, 1.48 mmol, 79%) as a white solid. **^1^H NMR** (300 MHz, MeOD) δ 4.67 – 4.54 (m, 1H), 3.91 (d, *J* = 4.7 Hz, 1H), 3.19 (q, *J* = 7.2 Hz, 6H), 1.48 (s, 9H), 1.45 (s, 9H), 1.38 – 1.26 (m, 12H). **^13^C NMR** (75 MHz, MeOD) δ 171.4, 158.2, 82.9, 80.6, 72.3, 61.9, 47.6, 28.7, 28.3, 19.5, 9.2. **^31^P NMR** (121 MHz, MeOD) δ 1.5. **HRMS** (ESI) m/z: [M + Na]^+^ Calcd for C_13_H_26_NO_8_PNa^+^ 378.1288; Found 378.1291.

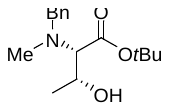
***N*-benzyl-*N*-methyl-l-threonine *tert*-butyl ester (S6)**

Compound **S6** was prepared according to literature procedure.^2^ l-threonine *tert*-butyl ester hydrochloride (2.23 g, 10.00 mmol, 1 eq.) was washed with NaHCO_3_ (sat. aq.). The aqueous layer was extracted with EtOAc (4x), the combined organic layers were washed with brine, dried over Na_2_SO_4_ and concentrated *in vacuo*. The residue was dissolved in dry methanol (100 mL, 0.1 M), after which benzaldehyde (1.07 mL, 10.50 mmol, 1.05 eq.) was added and the mixture was stirred at rt for 1 h. NaBH_3_(CN) (660 mg, 10.50 mmol, 1.05 eq.) was added and the mixture was stirred at rt for 1 h. When TLC analysis indicated full conversion, *para*-formaldehyde (905 mg, 31.50 mmol, 3.15 eq.) was added and the mixture was stirred for 4 h at rt NaBH_3_(CN) (660 mg, 10.50 mmol, 1.05 eq.) was added and the mixture was stirred at rt for 18 h. Once TLC analysis indicated conversion the reaction mixture was concentrated *in vacuo*, taken up in EtOAc and the resulting suspension was filtered through Celite. The resulting filtrate was concentrated *in vacuo* after which column chromatography (pentane/EtOAc/TEA, 99:1/1→97/3/1) yielded the desired product (1.35 g, 4.80 mmol, 48%) as a clear oil. **^1^H NMR** (400 MHz, CDCl_3_) δ 7.36 – 7.24 (m, 5H), 3.99 – 3.89 (m, 1H), 3.85 (d, *J* = 13.1 Hz, 1H), 3.62 (d, *J* = 13.0 Hz, 1H), 2.90 (d, *J* = 9.8 Hz, 1H), 2.32 (s, 3H), 1.53 (s, 9H), 1.17 (d, *J* = 6.0 Hz, 3H). **^13^C NMR** (101 MHz, CDCl_3_) δ 169.2, 138.3, 129.1, 128.6, 127.5, 81.8, 72.3, 63.0, 59.5, 37.8, 28.5, 19.3. **HRMS** (ESI) m/z: [M + H]^+^ Calcd for C_16_H_26_NO_3_^+^ 280.1907; Found 280.1872.

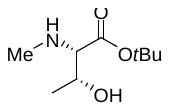
***N*-methyl-l-threonine *tert*-butyl ester (S7)**

Compound **S6** (4.02 g, 14.59 mmol, 1 eq.) was dissolved in methanol (144 mL, 0.1 M), the solution was purged with nitrogen gas and palladium on activated carbon (10 wt%, 766 mg, 0.72 mmol, 0.05 eq.) was added at rt. The solution was purged with nitrogen gas for 5 minutes, and then with hydrogen gas for 20 minutes. When TLC analysis indicated full conversion of starting material, the solution was purged with nitrogen gas for 5 minutes, filtered over Celite, and the filtrate was concentrated *in vacuo* to yield the desired product (2.47 g, 13.04 mmol, 89%) as clear oil. The product was used without further purification **^1^H NMR** (400 MHz, CDCl_3_) δ 3.64 – 3.52 (m, 1H), 3.21 (s, 1H), 2.74 (d, *J* = 7.8 Hz, 1H), 2.37 (s, 3H), 1.44 (s, 9H), 1.17 (d, *J* = 6.2 Hz, 3H). **^13^C NMR** (101 MHz, CDCl_3_) δ 172.7, 88.2, 82.0, 70.7, 67.8, 35.0, 28.1, 19.5.

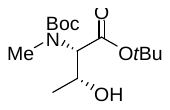
***N*-(tert-butoxycarbonyl)-*N*-methyl-l-threonine *tert*-butyl ester (S8)**

Compound **S7** (2.47 g, 13.04 mmol, 1 eq.) was dissolved in 1,4-dioxane:H2O (120 mL, 1:1, 0.1 M). Na_2_CO_3_ (2.76 g, 26.08 mmol, 2 eq.) and Boc2O (7.5 mL, 32.60 mmol, 2.5 eq.) were added and the mixture was stirred overnight at rt. When TLC analysis indicated full conversion, the mixture was washed with NaHCO3 (sat. aq.) and extracted with DCM (3x). The combined organic layers were dried over Na2SO4, filtered, and concentrated *in vacuo*. Column chromatography (pentane/EtOAc, 8/2) furnished the title compound (3.07 g, 10.76 mmol, 83%) as a clear oil. Note: Compound behaves as a rotamer in NMR. **^1^H NMR** (400 MHz, CDCl_3_) δ 4.37 – 4.26 (m, 2H), 3.18 (s, 1H), 2.88 (s, 3H), 1.42 (s, 18H), 1.16 (d, *J* = 5.8 Hz, 3H). **^13^C NMR** (101 MHz, CDCl_3_) δ 169.8, 157.1, 82.0, 80.3, 67.2, 65.2, 28.4, 28.1. **HRMS** (ESI) m/z: [M + H]^+^ Calcd for C_14_H_28_NO_5_^+^ 290.1962; Found 290.1959.

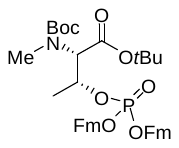
***N*-(tert-butoxycarbonyl)-*N*-methyl-*O*-(****bis((9H-fluoren-9-yl)methoxy))phosphoryl-l-threonine *tert*-butyl ester (S9)**

Threonine **S8** (289 mg, 1.00 mmol, 1 eq.) and phosphoramidite **S4** (574 mg, 1.10 mmol, 1.1 eq.) were co-evaporated with toluene (3x) before being dissolved in dry MeCN (12 ml, final concentration 0.05 M). Freshly activated 3Å molecular sieves were added and the mixture was allowed to stir 30 minutes at rt before adding DCI (0.25 M in MeCN; 8 mL, 2.00 mmol, 2 eq.). The reaction mixture was stirred for another 30 minutes until ^31^P NMR revealed full conversion. Upon completion, *t*BuOOH (5-6 M in decane, 0.55 mL, 3.00 mmol, 3 eq.) was added and the reaction mixture was stirred for 1 h until ^31^P NMR revealed full oxidation. NaHCO_3_ (sat. aq.) was added and the aqueous layer was extracted with EtOAc (3x), washed with brine (1x), dried over Na_2_SO_4_, filtered and concentrated *in vacuo*. Column chromatography (pentane/EtOAc/toluene, 80/20/2🡪60/40/2) provided phosphotriester **S9** (730 mg, 1.00 mmol, quantitative) as foamy solid. Note: Compound behaves as a rotamer in NMR. **^1^H NMR** (400 MHz, CDCl_3_) δ 7.76 – 7.66 (m, 4H), 7.53 (m, 3H), 7.47 – 7.27 (m, 7H), 7.25 – 7.18 (m, 2H), 5.02 – 4.90 (m, 1H), 4.80 (m, 1H), 4.43 (d, *J* = 4.6 Hz, 1H), 4.32 – 4.24 (m, 2H), 4.23 – 4.14 (m, 2H), 4.14 – 4.06 (m, 2H), 2.78 (2x s, 3H), 1.44 – 1.38 (3x s, 18H), 1.26 (2x d, *J* = 6.3 Hz, 3H). **^13^C NMR** (101 MHz, CDCl_3_) δ 167.9, 167.7, 156.7, 155.6, 143.3, 143.1, 143.1, 141.5, 141.4, 128.0, 128.0, 127.2, 127.2, 125.3, 125.2, 125.2, 125.1, 120.1, 120.1, 120.0, 82.5, 82.4, 80.7, 80.3, 75.2, 75.2, 74.2, 74.1, 69.3, 69.2, 69.1, 69.1, 64.1, 64.0, 62.3, 62.3, 48.0, 47.9, 47.9, 32.7, 32.3, 28.4, 28.4, 28.0, 19.1, 18.6. **^31^P NMR** (162 MHz, CDCl_3_) δ -2.3, -2.4. **HRMS** (ESI) m/z: [M + H]^+^ Calcd for C_42_H_49_NO_8_P^+^ 726.3190; Found 726.3146.

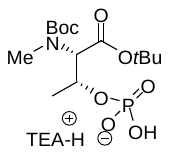
***N*-(tert-butoxycarbonyl)-*N*-methyl-*O*-phosphono-l-threonine *tert*-butyl ester triethylammonium salt (S10)**

Phosphotriester **S9** (2.13 g, 2.42 mmol, 1 eq.) was dissolved in MeCN (21 mL, 0.1 M) before adding TEA (5.1 mL, 36.3 mmol, 14 eq.). The reaction mixture was stirred overnight at rt until ^31^P NMR revealed full conversion to the monophosphate. The mixture was then concentrated *in vacuo* and column chromatography (DCM/MeOH, 100/0🡪90/10) furnished phosphate **S10** (911 mg, 1.94 mmol, 79%) as a white solid. Note: Compound behaves as a rotamer in NMR. **^1^H NMR** (400 MHz, MeOD) δ 4.87 – 4.74 (m, 1H), 4.55 (2x d, *J* = 6.4 Hz, 1H), 3.13 (q, *J* = 7.3 Hz, 4H), 3.00 (s, 3H), 1.52 – 1.43 (3x s, 18H), 1.40 – 1.35 (m, 3H), 1.29 (t, *J* = 7.3 Hz, 6H).**^13^C NMR** (101 MHz, MeOD) δ 170.3, 170.2, 158.3, 157.6, 83.0, 81.5, 81.2, 71.8, 71.7, 71.2, 71.2, 66.0, 65.9, 64.6, 64.6, 47.1, 33.3, 32.3, 28.7, 28.3, 19.7, 19.5, 9.1. **^31^P NMR** (162 MHz, MeOD) δ 0.2. **HRMS** (ESI) m/z: [M + Na]^+^ Calcd for C_14_H_28_NO_8_PNa^+^ 392.1445; Found 392.1445.

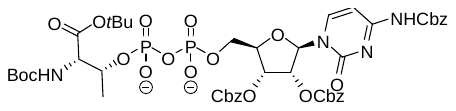
**Protected cytidine diphosphate l-threonine (S11)**

Phosphate **3** (228 mg, 0.50 mmol, 1 eq.) and amidite **5** (465 mg, 0.55 mmol, 1.1 eq.) were co-evaporated with MeCN (3x) before dissolving in MeCN (1.0 mL, final concentration 0.1 M). Freshly activated 3Å molecular sieves were added and the solution was stirred for 30 minutes before adding DCI (0.25 M in MeCN, 9 mL, 1.00 mmol, 2 eq.). The reaction mixture was stirred for 30 minutes until ^31^P NMR revealed full conversion. Upon completion, *t*BuOOH (5-6 M in decane, 0.18 mL, 1.00 mmol, 2 eq.) was added and the reaction mixture was stirred for 1 h. When ^31^P NMR revealed full oxidation, DBU (0.75 mL, 5.00 mmol, 10 eq.) was added and the mixture was stirred for 20 minutes until ^31^P NMR revealed full deprotection. The reaction mixture was quenched by the addition of pyridinium chloride (578 mg, 5.00 mmol, 10 eq.), stirred for 10 minutes and concentrated *in vacuo*. Size exclusion chromatography (LH-20, DCM/MeOH) provided the crude pyrophosphate (560 mg). The product was used without further purification.

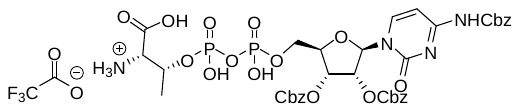
***N*^4^-benzyloxycarbonyl-2’,3’-di-*O*-(benzyloxycarbonyl) cytidine diphosphate l-threonine** (**6**)

Pyrophosphate **S11** (560 mg, ~0.41 mmol, 1 eq.) was dissolved in DCM (8.2 mL, 0.05 M) and cooled to 0°C. TFA (8.2 mL, 50v/v%) was added and the reaction mixture was stirred for 5 h until LC-MS revealed deprotection of the *tert*-butyl ester and BOC groups. The mixture was concentrated *in vacuo*, co-evaporated with DCM (3x) and toluene (3x). HPLC purification (C18 column, H_2_O/MeCN/TFA method) provided the partly deprotected target compound (69 mg, 73 µmol, 18% over 4 steps) as a white solid. **^1^H NMR** (400 MHz, MeOD) δ 8.41 (d, *J* = 7.6 Hz, 1H), 7.41 – 7.25 (m, 16H), 6.18 (d, *J* = 5.1 Hz, 1H), 5.49 (t, *J* = 4.7 Hz, 1H), 5.44 (t, *J* = 5.2 Hz, 1H), 5.19 (s, 2H), 5.16 – 5.00 (m, 4H), 4.64 – 4.54 (m, 1H), 4.45 – 4.39 (m, 1H), 4.36 – 4.18 (m, 2H), 3.69 (d, *J* = 8.7 Hz, 1H), 1.52 (d, *J* = 6.2 Hz, 3H). **^13^C NMR** (101 MHz, MeOD) δ 165.29, 155.40, 155.26, 146.37, 129.59, 129.55, 129.52, 129.44, 129.39, 97.97, 89.19, 83.03, 78.67, 75.45, 72.81, 71.37, 71.18, 68.58, 65.79, 60.97, 20.19. **^31^P NMR** (202 MHz, MeOD) δ -9.5, -9.6, -9.9, -10.0. **HRMS** (ESI) m/z: [M + H]^+^ Calcd for C_37_H_41_N_4_O_19_P_2_^+^ 907.1835; Found 907.1832.

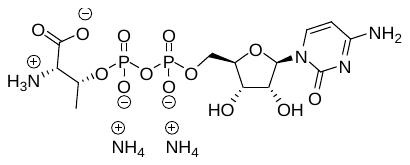
**Cytidine diphosphate l-threonine (CDP-Thr)**

The hydrogen transfer was performed according to literature.^3^ Compound **6** (6 mg, 6 µmol, 1 eq.) was dissolved in EtOH/H_2_O (50v/v%, 0.6 mL, 0.01 M) and Pd/C (10%wt, 16 mg, 0.015 mmol, 2.5 eq.) was added. 1,4-cyclohexadiene (28 µL, 300 µmol, 50 eq.) was added and the reaction mixture was stirred for 2 h until LC-MS revealed full reduction. The mixture was over Celite, lyophilized and gel filtration (HW-40, 0.15 M NH_4_OAc (aq.) with 10% MeCN) provided **CDP-Thr** (1.7 mg, 3.2 µmol, 50%) as a white solid. **^1^H NMR** (500 MHz, D_2_O) δ 7.97 (d, *J* = 7.6 Hz, 1H), 6.13 (d, *J* = 7.7 Hz, 1H), 5.97 (d, *J* = 4.2 Hz, 1H), 4.67 – 4.55 (m, 1H), 4.33 (t, *J* = 5.0 Hz, 1H), 4.30 (d, *J* = 5.0 Hz, 1H), 4.28 – 4.23 (m, 2H), 4.21 – 4.14 (m, 1H), 3.66 (d, *J* = 7.1 Hz, 1H), 1.47 (d, *J* = 7.1 Hz, 3H). **^13^C NMR** (101 MHz, D_2_O) δ 171.71, 162.75, 142.75, 95.94, 89.41, 82.99, 82.90, 74.29, 72.31, 72.25, 69.16, 64.60, 64.55, 60.35, 60.27, 18.68. **^31^P NMR** (202 MHz, D_2_O) δ -10.3, -10.4, -11.7, -11.8. **HRMS** (ESI) m/z: [M + H]^+^ Calcd for C_37_H_41_N_4_O_19_P_2_^+^ 505.0731; Found 505.0738.

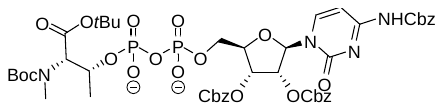
**Protected cytidine diphosphate *N*-methyl l-threonine (S11)**

Phosphate **S10** (74 mg, 0.20 mmol, 1 eq.) and amidite **5** (170 mg, 0.22 mmol, 1.1 eq.) were co-evaporated with MeCN (3x) before dissolving in MeCN (0.5 mL, final concentration 0.1 M). Freshly activated 3Å molecular sieves were added and the solution was stirred for 30 minutes before adding DCI (0.25 M in MeCN, 1.6 mL, 0.40 mmol, 2 eq.). The reaction mixture was stirred for 30 minutes until ^31^P NMR revealed full conversion. Upon completion, *t*BuOOH (5-6 M in decane, 0.15 mL, 0.80 mmol, 4 eq.) was added and the reaction mixture was stirred for 1 h. When ^31^P NMR revealed full oxidation, DBU (0.30 mL, 2.00 mmol, 10 eq.) was added and the mixture was stirred for 20 minutes until ^31^P NMR revealed full deprotection. The reaction mixture was quenched by the addition of pyridinium chloride (231 mg, 2.00 mmol, 10 eq.), stirred for 10 minutes and concentrated *in vacuo*. Size exclusion chromatography (LH-20, DCM/MeOH) provided the crude pyrophosphate (207 mg). The product was used without further purification.

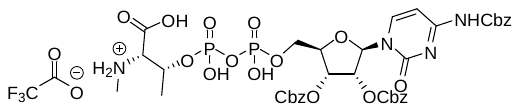
***N*^4^-benzyloxycarbonyl-2’,3’-di-*O*-(benzyloxycarbonyl) cytidine diphosphate *N*-methyl-l-threonine** (**S12**)

Pyrophosphate **S11** (207 mg, ~0.14 mmol, 1 eq.) was dissolved in DCM (2.88 mL, 0.05 M) and cooled to 0°C. TFA (1.44 mL, 50v/v%) was added and the reaction mixture was stirred for 5 h until LC-MS revealed deprotection of the *tert*-butyl ester and BOC groups. The mixture was concentrated *in vacuo*, co-evaporated with DCM (3x) and toluene (3x). HPLC purification (C18 column, H_2_O/MeCN/TFA method) provided the partly deprotected target compound (42 mg, 45 µmol, 31% over 4 steps) as a white solid. **^1^H NMR** (400 MHz, MeOD) δ 8.42 (d, *J* = 7.6 Hz, 1H), 7.44 – 7.26 (m, 16H), 6.20 (d, *J* = 4.8 Hz, 1H), 5.55 – 5.47 (m, 2H), 5.22 (s, 2H), 5.18 – 5.03 (m, 4H), 4.61 – 4.50 (m, 1H), 4.49 – 4.40 (m, 1H), 4.40 – 4.20 (m, 2H), 3.40 (d, *J* = 9.3 Hz, 1H), 2.66 (s, 3H), 1.51 (d, *J* = 6.3 Hz, 3H). **^13^C NMR** (101 MHz, MeOD) δ 170.8, 165.2, 157.8, 155.4, 155.3, 154.3, 146.4, 137.2, 136.6, 136.4, 129.6, 129.4, 129.4, 97.9, 89.5, 82.9, 82.8, 78.6, 75.3, 73.3, 73.2, 71.8, 71.8, 71.4, 71.2, 68.6, 65.8, 65.8, 33.1, 20.2. **^31^P NMR** (162 MHz, MeOD) δ -10.5, -10.6, -11.3, -11.4. **HRMS** (ESI) m/z: [M + H]^+^ Calcd for C_38_H_43_N_4_O_19_P_2_^+^ 921.1991; Found 921.1992.

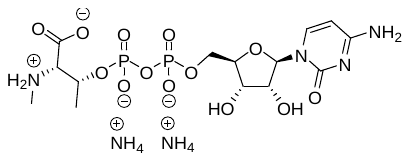
**Cytidine diphosphate *N*-methyl l-threonine (CDP-MeThr)**

The hydrogen transfer was performed according to literature.^3^ Compound **S12** (12 mg, 13 µmol, 1 eq.) was dissolved in EtOH/H_2_O (50v/v%, 1.3 mL, 0.01 M) and Pd/C (10%wt, 32 mg, 0.030 mmol, 2.5 eq.) was added. 1,4-cyclohexadiene (57 µL, 600 µmol, 50 eq.) was added and the reaction mixture was stirred overnight until LC-MS revealed full reduction. The mixture was over Celite, lyophilized and gel filtration (HW-40, 0.15 M AcOH (aq.) with 10% MeCN) provided **CDP-MeThr** (1.1 mg, 1.9 µmol, 9%) as a white solid. **^1^H NMR** (500 MHz, D_2_O) δ 7.94 (d, *J* = 7.6 Hz, 1H), 6.09 (d, *J* = 7.7 Hz, 1H), 5.90 (d, *J* = 3.9 Hz, 1H), 4.48 – 4.36 (m, 1H), 4.29 – 4.18 (m, 4H), 4.16 – 4.08 (m, 1H), 3.44 (d, *J* = 8.8 Hz, 1H), 2.64 (s, 3H), 1.38 (d, *J* = 6.3 Hz, 3H). **^31^P NMR** (202 MHz, D_2_O) δ -10.6, -10.7, -12.2, -12.3. **HRMS** (ESI) m/z: [M + H]^+^ Calcd for C_14_H_25_N_4_O_13_P_2_^+^ 519.0888; Found 519.0888.

(1)	Bialy, L.; Waldmann, H. Total Synthesis and Biological Evaluation of the Protein Phosphatase 2A Inhibitor Cytostatin and Analogues. *Chemistry – A European Journal* **2004**, *10* (11), 2759–2780. https://doi.org/https://doi.org/10.1002/chem.200305543.

(2)	White, K. N.; Konopelski, J. P. Facile Synthesis of Highly Functionalized N-Methyl Amino Acid Esters without Side-Chain Protection. *Org. Lett.* **2005**, *7* (19), 4111–4112. https://doi.org/10.1021/ol051441w.

(3)	Johnson, D. C.; Widlanski, T. S. Facile Deprotection of O-Cbz-Protected Nucleosides by Hydrogenolysis:  An Alternative to O-Benzyl Ether-Protected Nucleosides. *Org. Lett.* **2004**, *6* (25), 4643–4646. https://doi.org/10.1021/ol048426w.

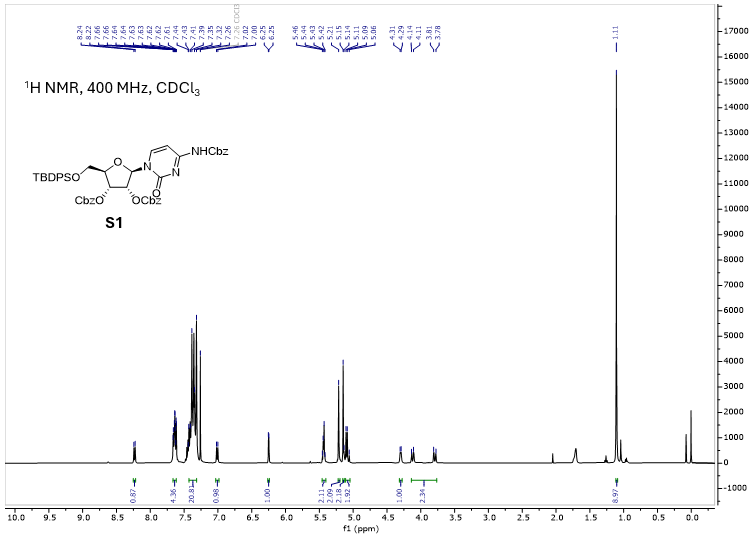

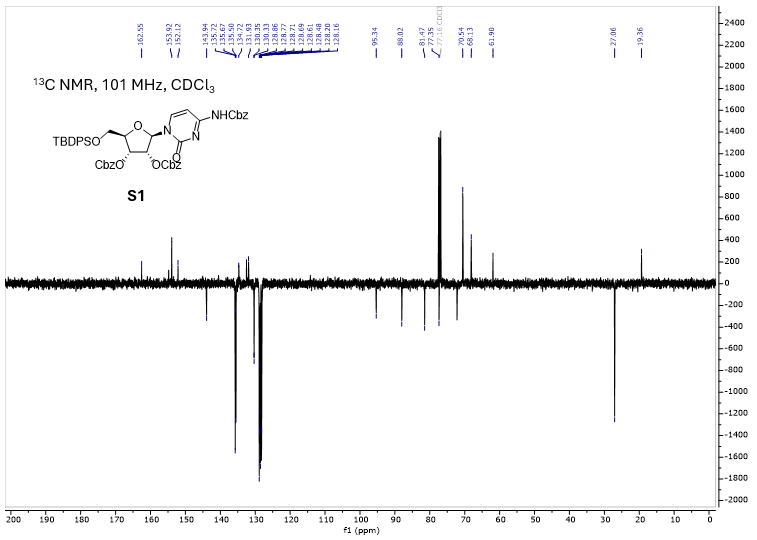

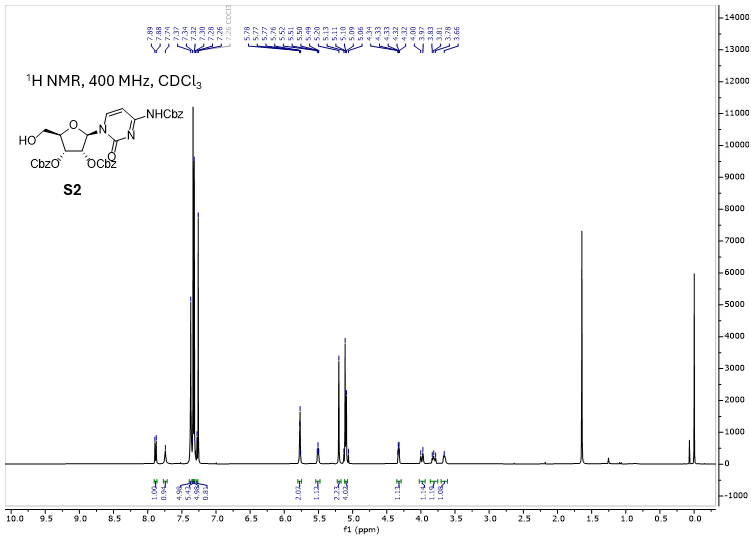

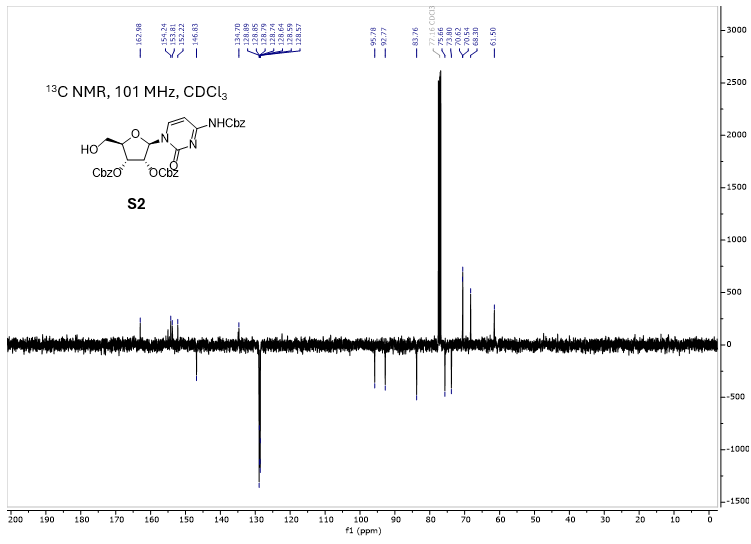

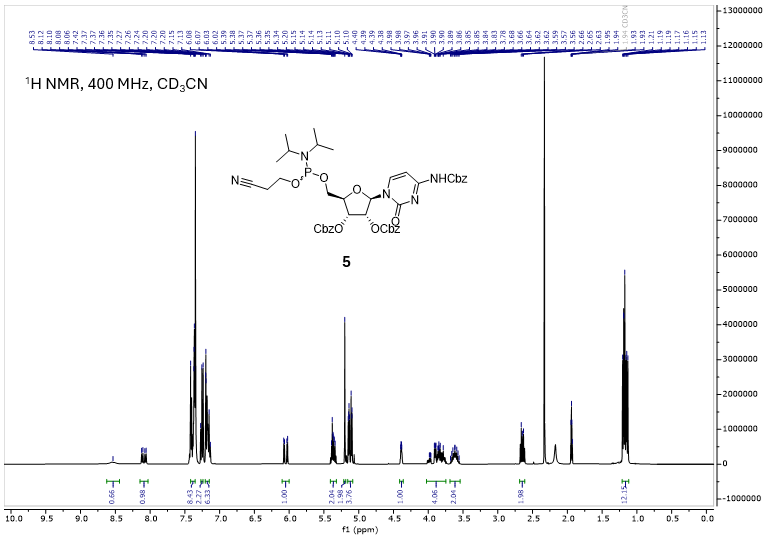

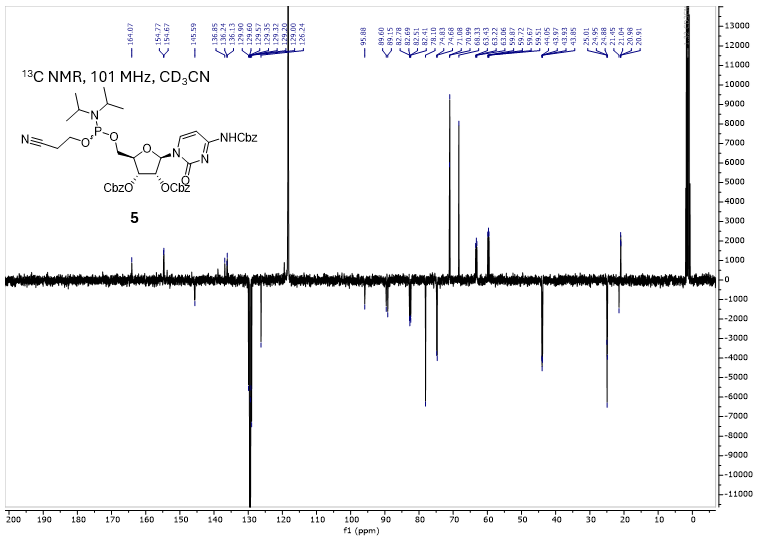

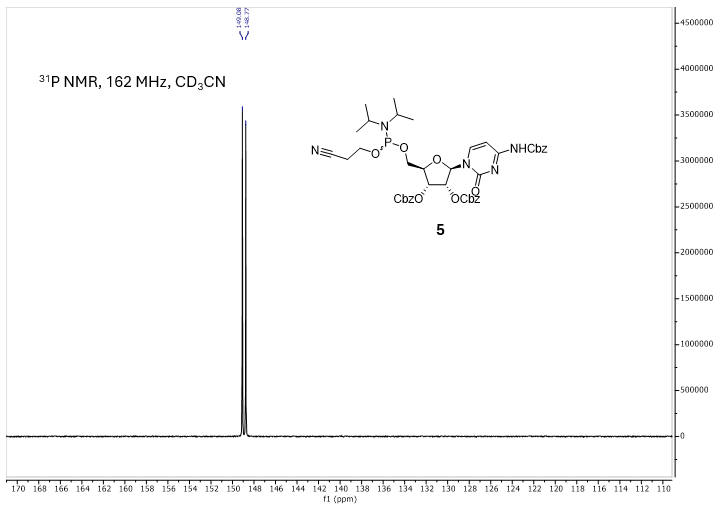

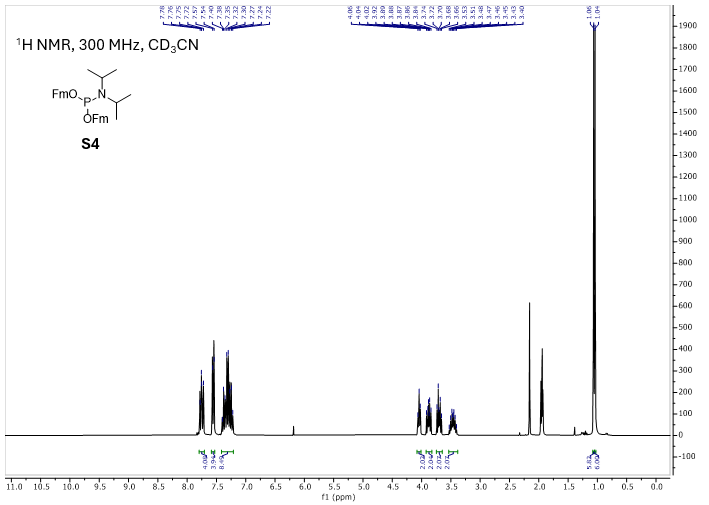

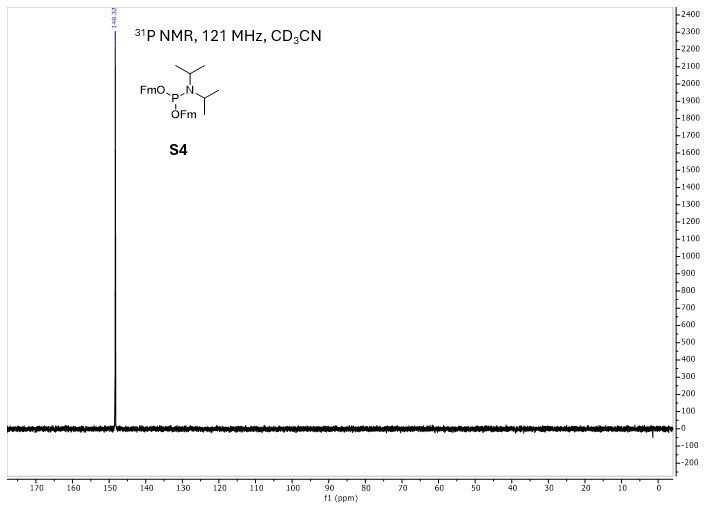

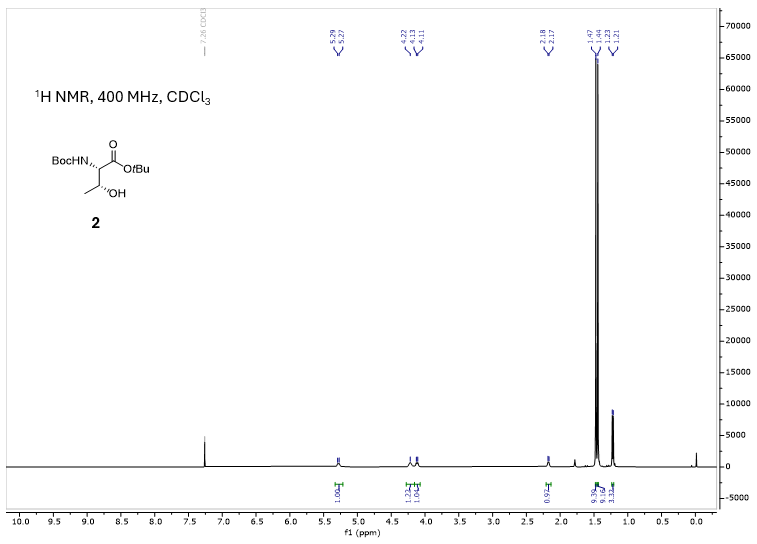

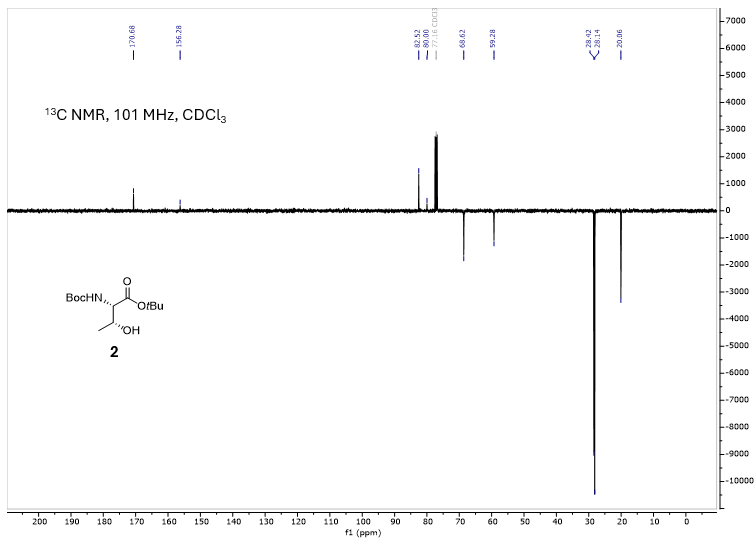

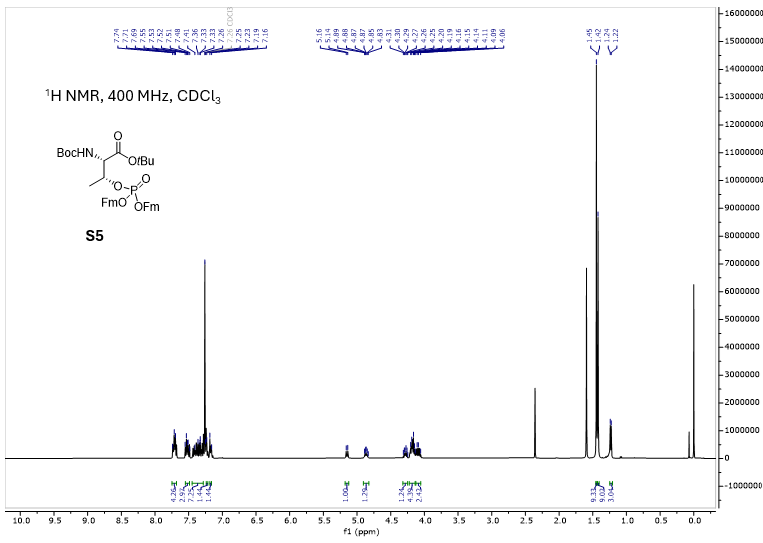

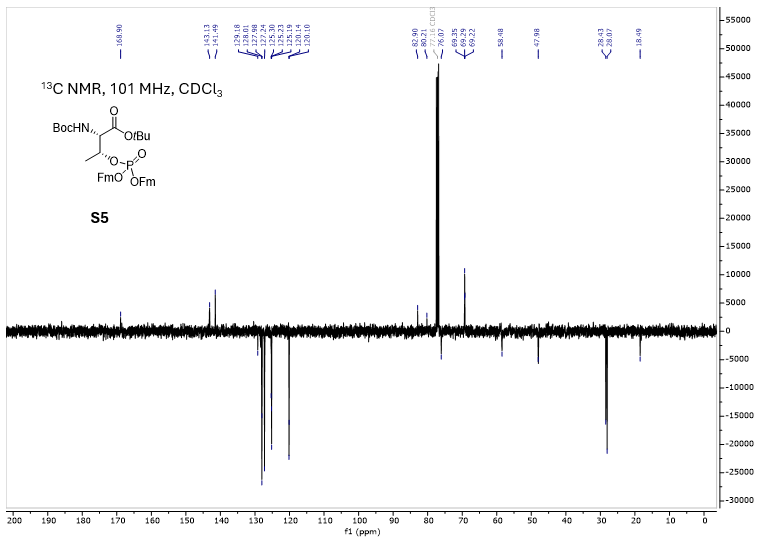

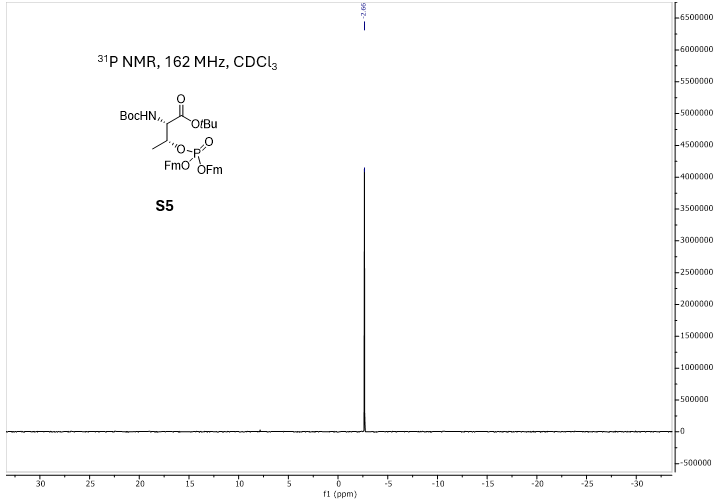

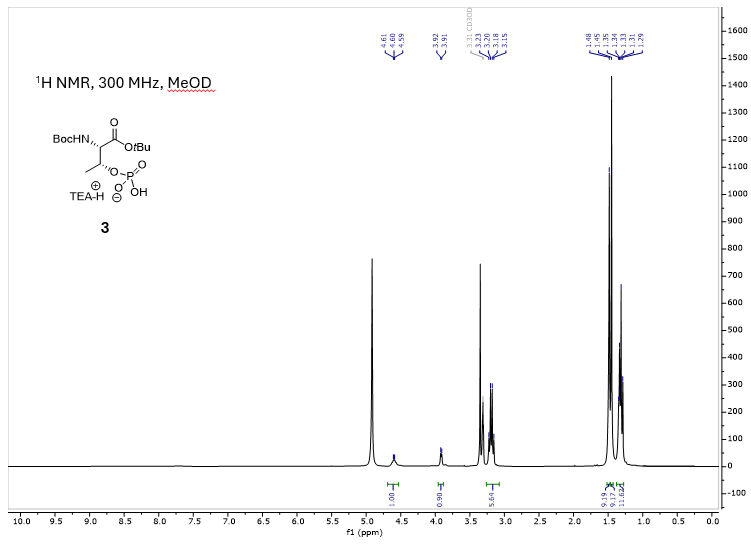

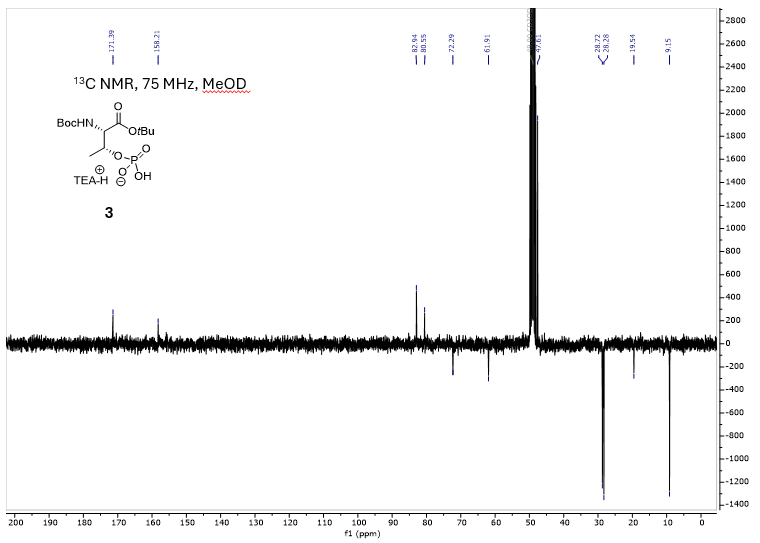

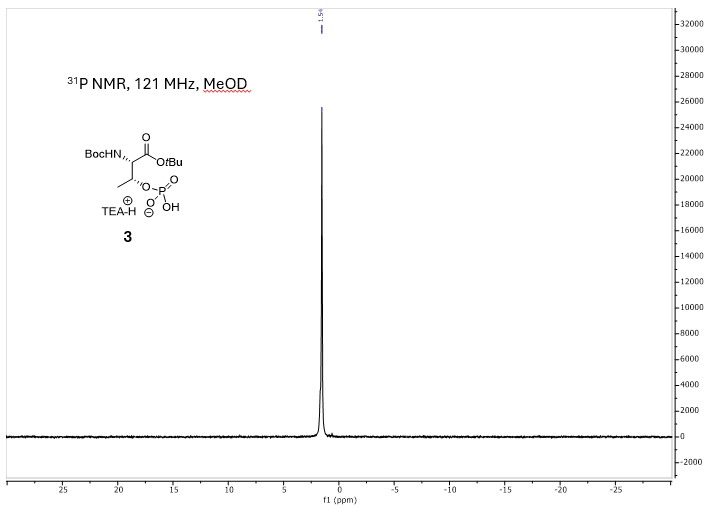

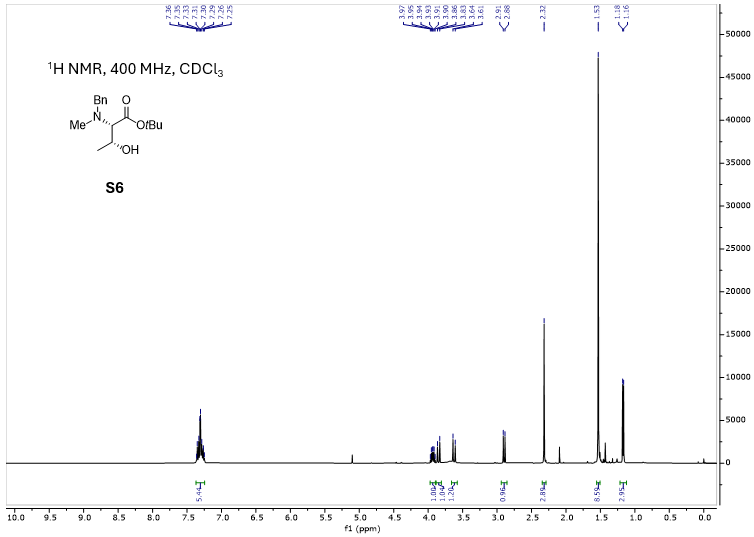

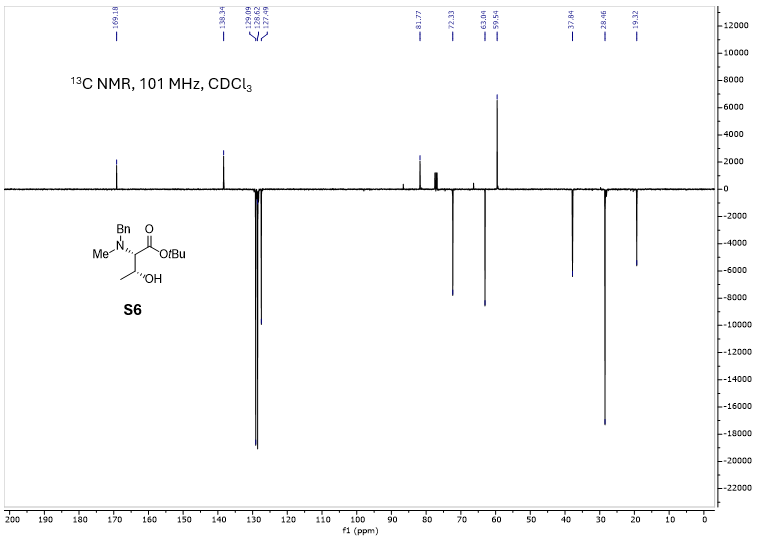

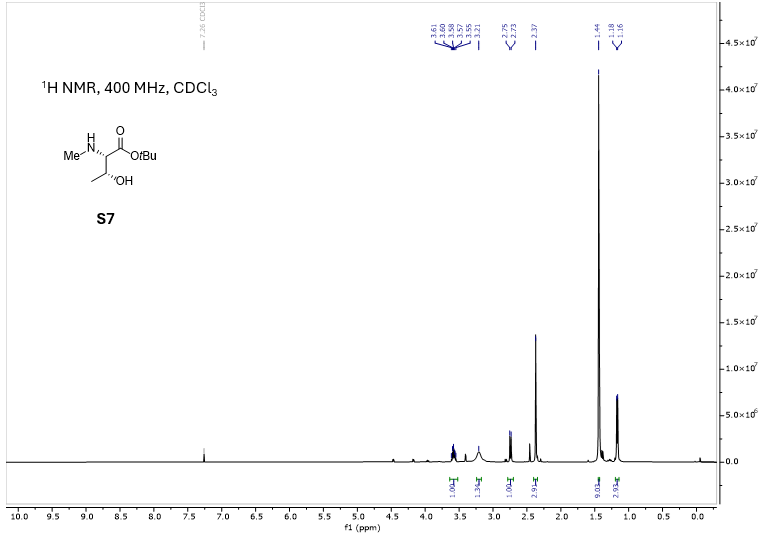

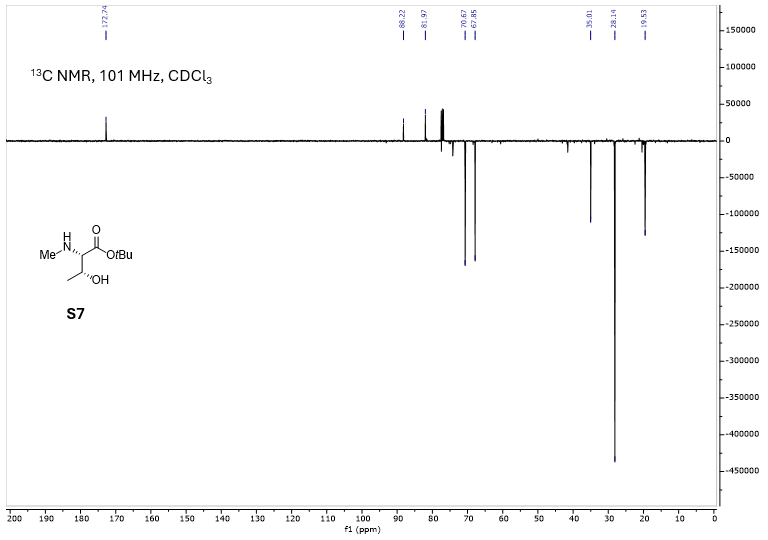

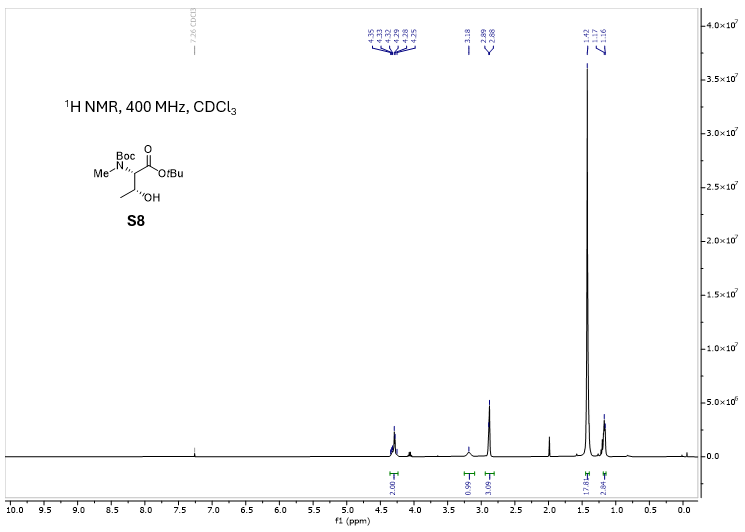

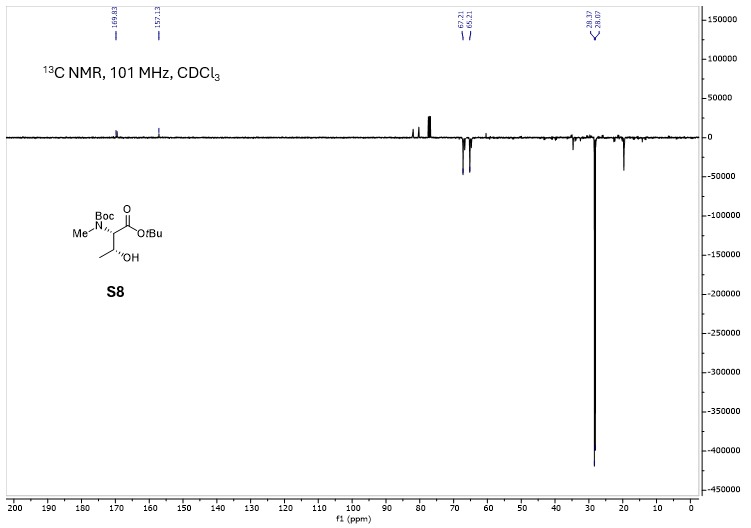

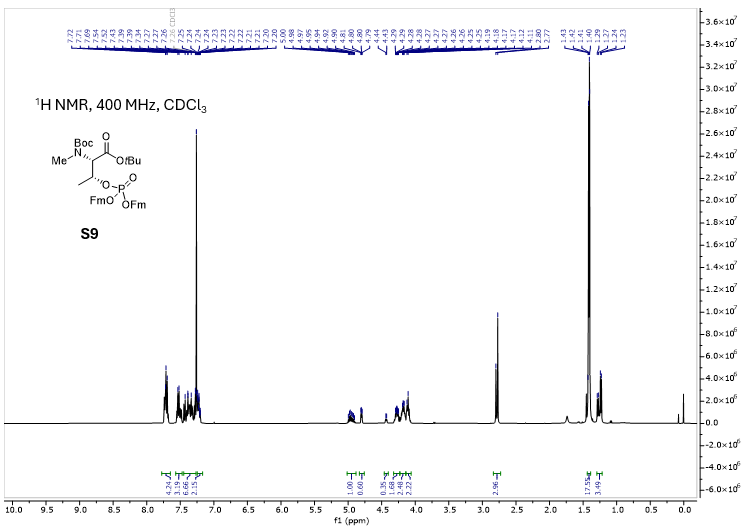

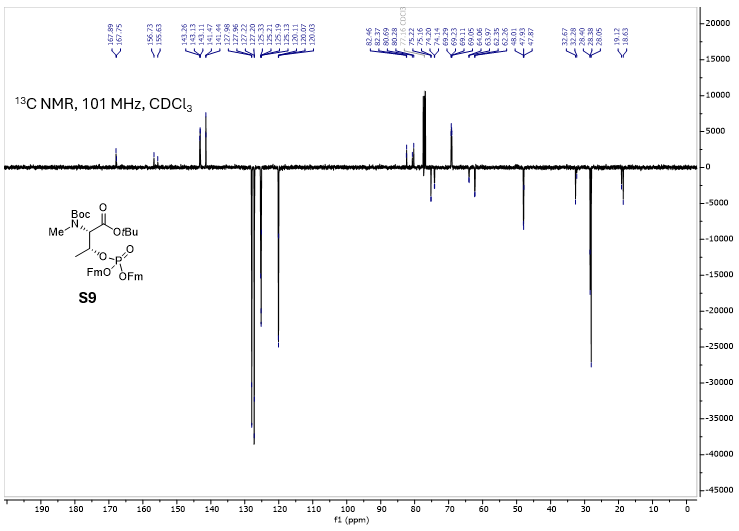

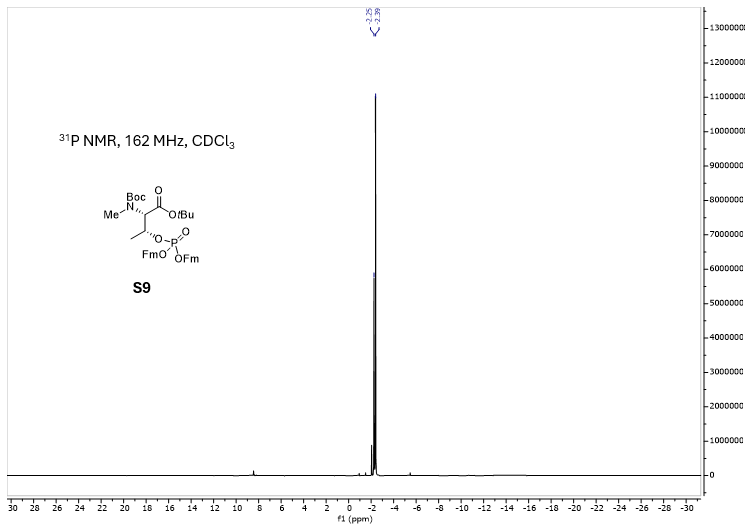

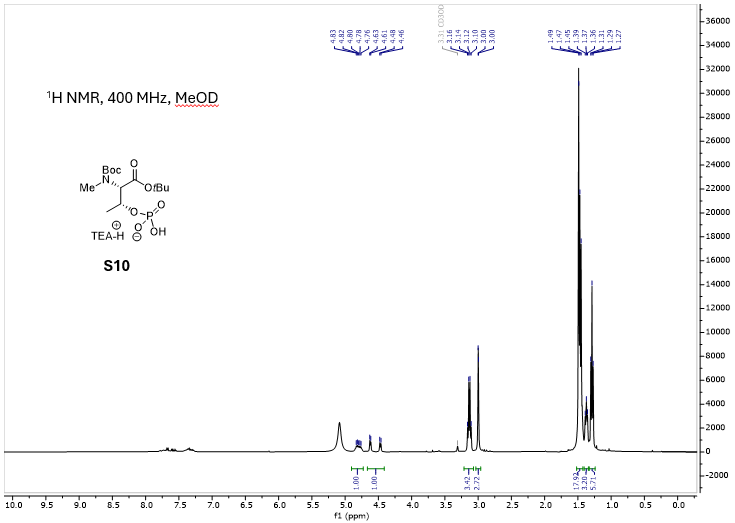

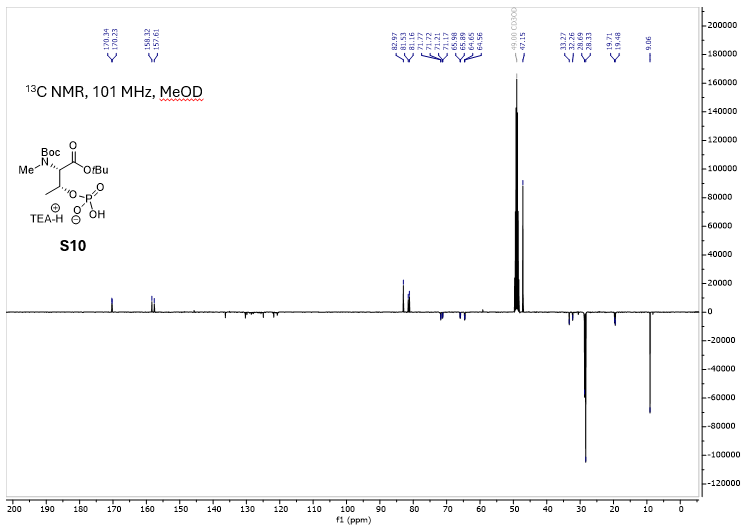

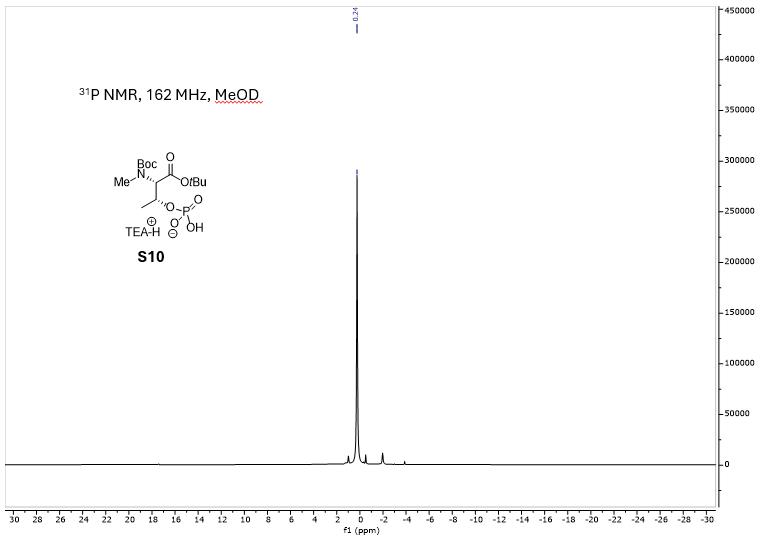

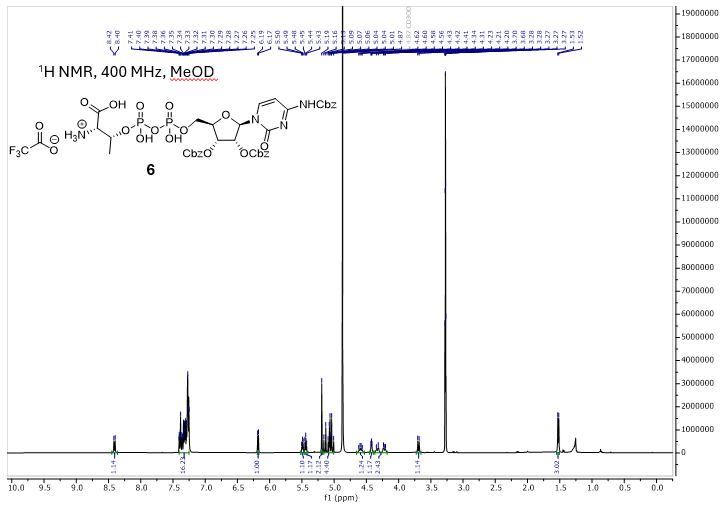

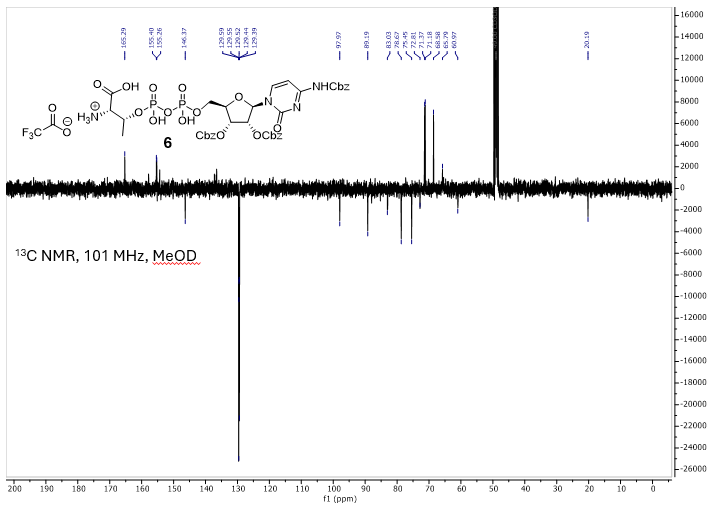

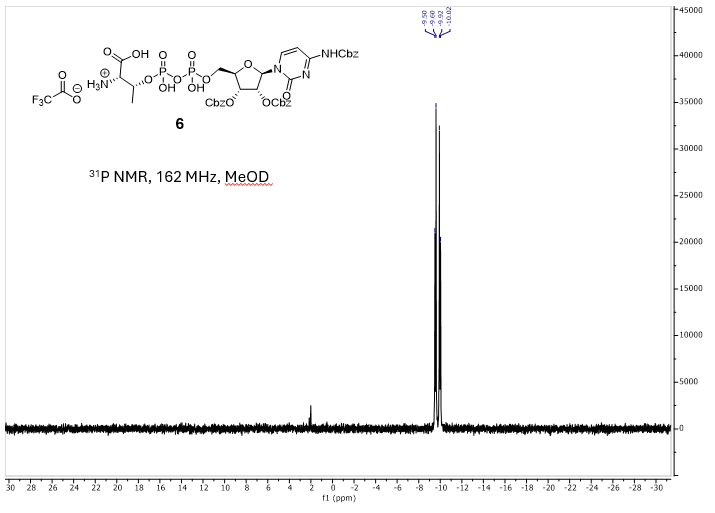

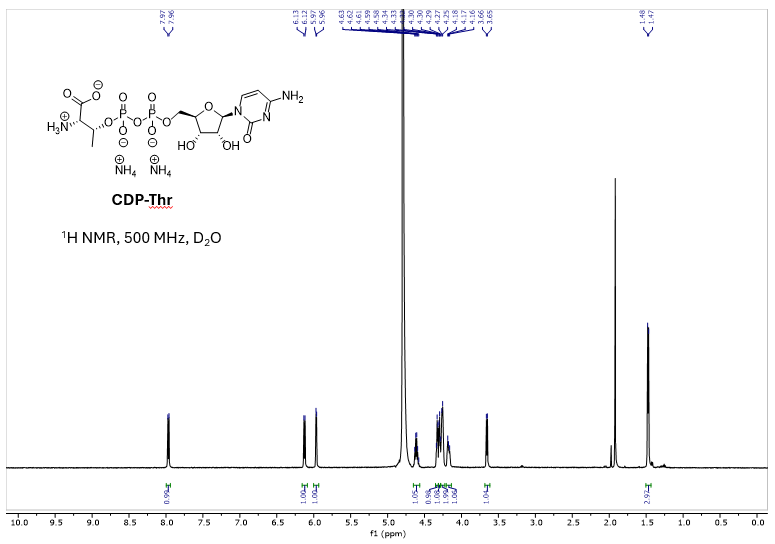

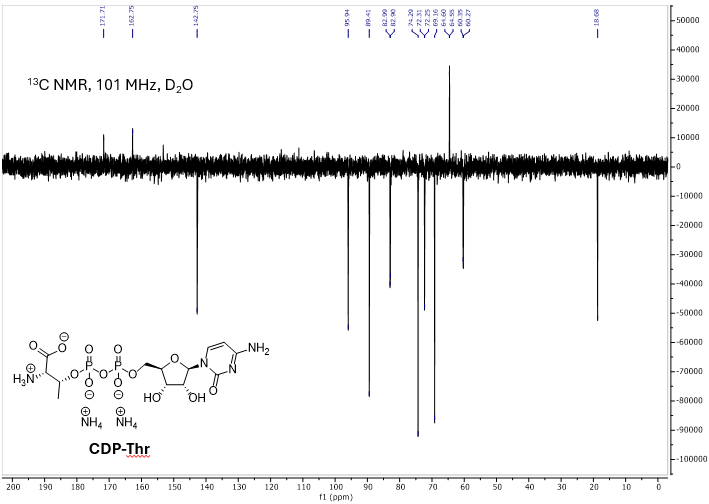

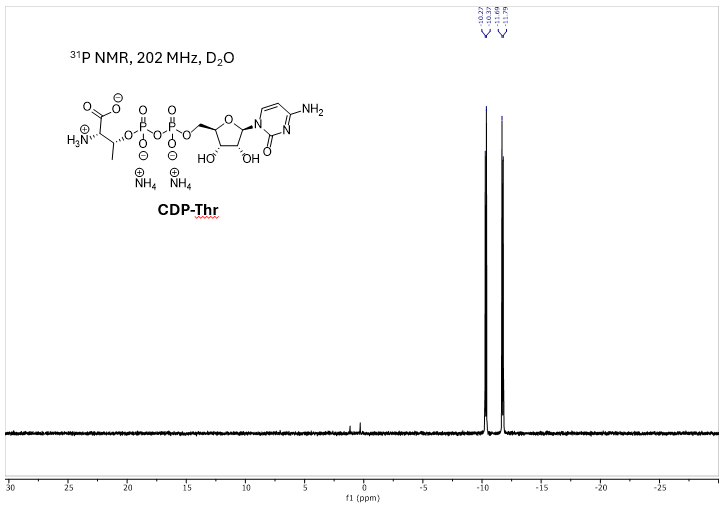

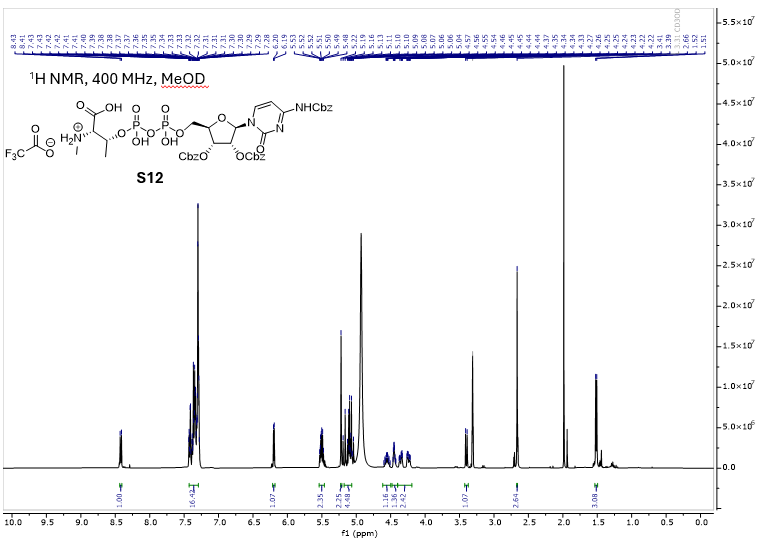

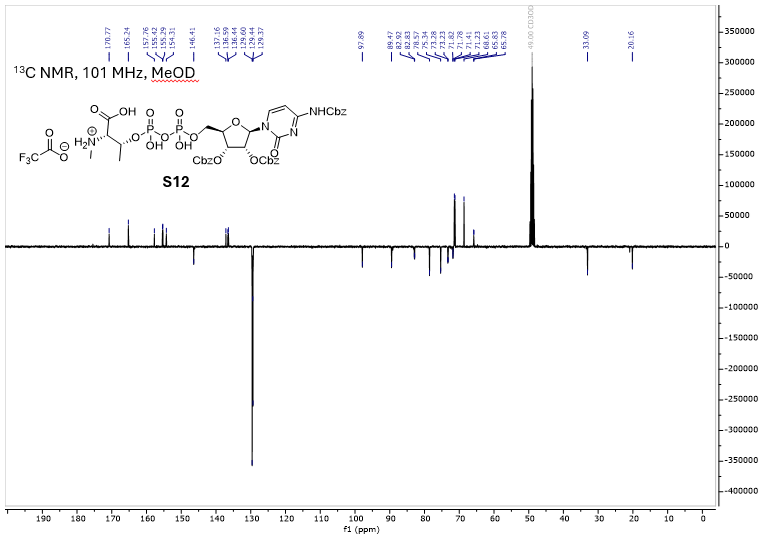

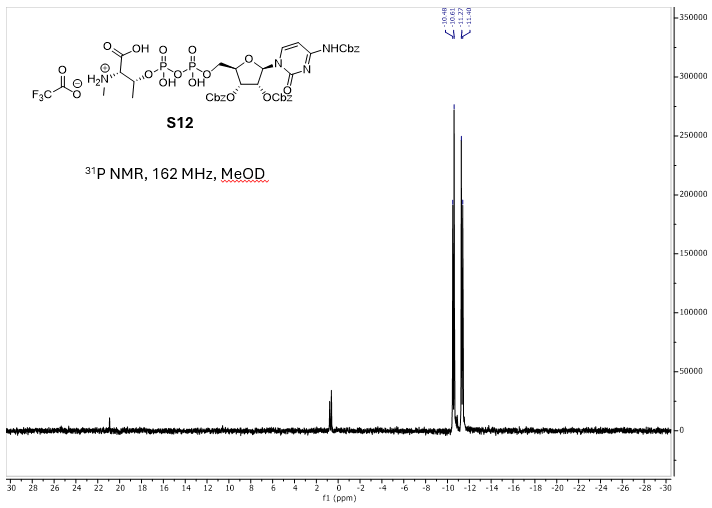

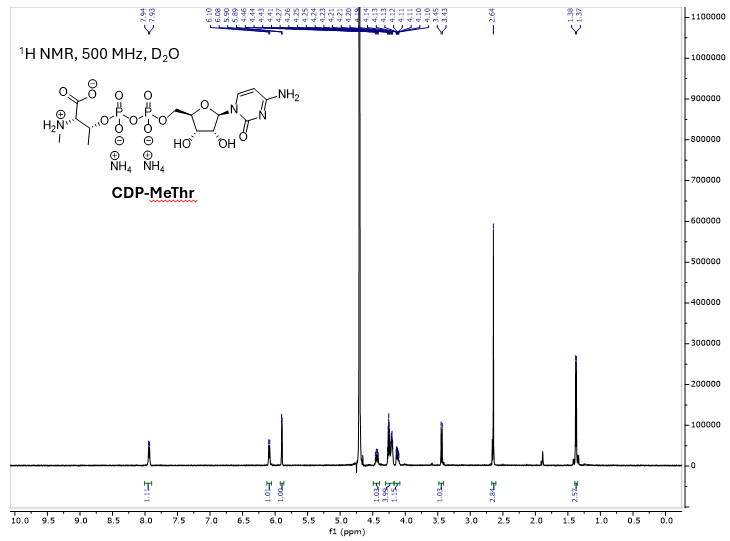

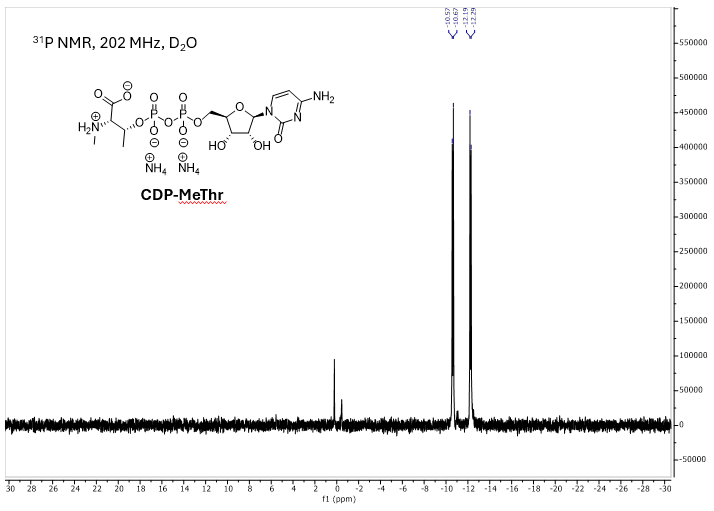
